# Maturation of human induced pluripotent stem cell-derived cardiomyocytes for modeling hypertrophic cardiomyopathy

**DOI:** 10.1101/2020.01.16.909598

**Authors:** Walter E. Knight, Yingqiong Cao, Ying-Hsi Lin, Genevieve C. Sparagna, Betty Bai, Yuanbiao Zhao, Congwu Chi, Yanmei Du, Pilar Londono, Julie A. Reisz, Benjamin C. Brown, Matthew R. G. Taylor, Amrut V Ambardekar, Joseph C. Cleveland, Timothy A. McKinsey, Mark Y. Jeong, Lori A. Walker, Kathleen C. Woulfe, Angelo D’Alessandro, Kathryn C. Chatfield, Hongyan Xu, Michael R. Bristow, Peter M. Buttrick, Kunhua Song

## Abstract

**Rationale:** Human induced pluripotent stem cell derived cardiomyocytes (hiPSC-CMs) are a powerful platform for biomedical research. However, they are immature, which is a barrier to modeling adult-onset cardiovascular disease.

**Objective:** We sought to develop a simple method which could drive cultured hiPSC-CMs towards maturity across a number of phenotypes.

**Methods and results:** Cells were cultured in fatty acid-based media and plated on micropatterned surfaces to promote alignment and elongation. These cells display many characteristics of adult human cardiomyocytes, including elongated cell morphology, enhanced maturity of sarcomeric structures, metabolic behavior, and increased myofibril contractile force. Most notably, hiPSC-CMs cultured under optimal maturity-inducing conditions recapitulate the pathological hypertrophy caused by either a pro-hypertrophic agent or genetic mutations.

**Conclusions:** The more mature hiPSC-CMs produced by the methods described here will serve as a useful *in vitro* platform for characterizing cardiovascular disease.

## Introduction

Human induced pluripotent stem cell derived cardiomyocytes (hiPSC-CMs) represent a powerful tool to study cardiovascular physiology and disease, offering a number of unique benefits. Significant differences exist between the hearts of humans and small animals most often used for modeling cardiovascular disease, including heart rate, cell size, multinucleation frequency, and myosin heavy chain expression ^1^. Therefore, the capability to study cardiovascular disease in a human cell system is critical. hiPSC-CMs offer a number of additional benefits, including scalability, the ability to generate patient-specific cells, and quickly create knockout and transgenic lines via CRISPR/Cas9-mediated gene editing. In spite of these benefits, hiPSC-CMs possess a significant drawback-they are functionally immature, and typically resemble fetal or neonatal cardiomyocytes in terms of cell size and morphology, gene expression, myofibril contractility, and metabolic activity^1–4^. While certain forms of cardiovascular disease, such as congenital heart defects and cardiomyopathies caused by homozygous or compound heterozygous mutations, do primarily affect infants, most forms of pathological cardiac remodeling are diseases that afflict adults. Therefore, the functional immaturity of hiPSC-CMs represents a potential barrier to modeling and investigating many forms of cardiovascular disease.

Here, we sought to develop a method to produce structurally and functionally mature hiPSC-CMs, which could be performed with relatively little specialized equipment and could be scaled up easily. To this end, we combined and optimized existing approaches which have previously been reported to induce hiPSC-CM maturation. First, we attempted to facilitate metabolic maturation by culturing our cells in maturation medium where fatty acids, rather than glucose, represent the primary energy source. Additionally, we cultured the cells on micropatterned surfaces, designed to promote cellular elongation. hiPSC-CMs were maintained in either standard glucose-based media (RPMI 1640, GLUC), with maturation medium (MM), or with maturation medium and patterning (MPAT). We found that compared to GLUC, both MM and MPAT hiPSC-CMs displayed a number of structural and functional improvements, including increased expression of fatty acid oxidizing genes, more mature mitochondria, and increased myofibril active tension generation. Additionally, MM and MPAT cells displayed a robust response to the hypertrophic agonist phenylephrine (PE), which could be inhibited by the bromodomain and extra-terminal domain (BET) inhibitor, JQ1. GLUC cells failed to respond to PE, potentially indicating that prolonged culture in glucose-containing medium induces a hypertrophic state. hiPSC-CMs derived from patients with Danon disease cultured in MM or MPAT cells also displayed a robust, spontaneous hypertrophic response compared to GLUC. Our results indicate that the combinatorial approach employed in MPAT culture produces hiPSC-CMs which demonstrate both adult-like myofibril mechanics and an adult-like hypertrophic response. The use of this approach will facilitate the study of adult cardiovascular disease with hiPSC-CMs.

## Methods and Materials

### Condensed Methods

An extended version of the methods section with more detailed protocols is available in the supplement.

### Human and Animal Subject Information

The portion of this study related to human subjects reviewed and approved by the Colorado Multiple Institutions Review Board (COMIRB #06-0452). Human hearts from healthy donors patients with hypertrophic cardiomyopathy (HCM) were obtained from the tissue bank maintained by the Division of Cardiology at the University of Colorado (COMIRB #01-568). All patients were followed by the University of Colorado Heart Failure Program and offered participation in the research protocol. All research involving animals complied with protocols approved by the Institutional Animal Care and Use Committee (IACUC) of University of Colorado.

### hiPSC-CM differentiation and culture

hiPSCs were induced into cardiomyocytes via modulation of Wnt signaling, according to previously established methods^5^. 20-25 days after induction, cardiomyocytes were purified using lactate medium^6^. hiPSC-CMs were cultured in RPMI-1640 (with 10.9mmol glucose) medium supplemented with B-27™ diluted 1:50 (GLUC) until approximately day 30 post induction; MM and MPAT cells were then maintained in fatty acid/galactose-based maturation medium (glucose-free RPMI, 50μmol palmitic/100μmol oleic acid, 10mmol galactose, and B-27™ diluted 1:50). hiPSC-CMs were replated (either onto fibronectin-coated unpatterned or patterned surfaces or standard cell culture dishes) approximately day 40 post induction.

### Patterned surface preparation

Patterned surfaces were prepared from clear plastic coverslips using 20 micrometer lapping paper, and sterilized.

### Myofibril mechanics

hiPSC-CMs were lysed using a cell scraper with a 20% sucrose in a relaxing solution. Myofibrils were mounted between two glass instruments, and contraction induced via introduction of calcium, followed by rapid calcium withdrawal via a pipette switching technique.

### CPT and Cardiolipin Assays

CPT and cardiolipin assays were performed on flash frozen cell pellets. CPT activity was assessed using a 14C radioactive assay. Cardiolipin content was assessed using mass spectrometry.

## Results

### Maturation methods improve both hiPSC-CM and sarcomeric morphology

To investigate whether hiPSC-CMs could be shifted towards more adult-like morphology and behavior, a combinatorial approach was employed. First, instead of using media where glucose is the main energy source (GLUC) our cells were cultured in media where glucose was substituted with a combination of galactose, oleic acid, and palmitic acid (maturity medium, MM) based on previous reports of similar methods that improve cellular morphology, contractility, ATP content, and metabolic behavior^7, 8^. Although both GLUC and MM each contain the B-27™ supplement, which contains several fatty acids, total fatty acid content in MM is ~150μmol versus ~144nM in B-27™-supplemented RPMI^9^. As an additional step to improve hiPSC-CM maturity, MM-cultured hiPSC-CMs were plated onto plastic coverslips which had been micropatterned with grooves via 20 micron lapping paper, to induce cell elongation and anisotropic alignment (MPAT). When plated onto these patterned surfaces, cells adhered within hours, grew into the grooves within 24-48 hours (**Fig. S1**) and displayed coordinated uniaxial contraction along the direction of patterning, which was not observed in other conditions (**Supplementary Video 1-3**).

We then assessed whether either patterning or MM induced changes in cellular morphology or sarcomere organization by immunofluorescent microscopy. Cells cultured under each condition expressed the sarcomeric proteins cardiac troponin (cTnI), alpha actinin, ventricular myosin light chain 2 (MLC-2V), and myosin binding protein C3 (MYBPC3) (**Fig. 1A**). In MM and MPAT cells, sarcomeres showed greater levels of organization, with Z-lines oriented perpendicularly to the long axis of the cells, whereas in GLUC, sarcomeres wrapped around the cell in a circular fashion, or were oriented chaotically throughout the cytosol. Additionally, some GLUC cells lacked sarcomeres in significant areas of cytosol (**Fig. 1A**, white arrows). We also assessed differences in cell morphology quantitatively. Cells cultured with MM or MPAT were approximately 30% smaller than cells cultured in GLUC, consistent with previous reports^7^ (**Fig. 1B**). MM cells also showed a significantly lower circularity value than GLUC cells. Circularity was further reduced in MPAT cells, indicating that MPAT cells were significantly more elongated than MM cells, given their similar size (**Fig. 1C**). While average cell area of GLUC cells was comparable to that of adult mouse cardiomyocytes, MM and MPAT cells were both significantly smaller, and all hiPSC-CM groups showed a significantly greater variability of cell area compared to adult mouse cells (**Fig. S2A**). All hiPSC-CM groups had cell areas lower than those reported for adult human cardiomyocytes (10-14,000μm^2^)^2^. More than 75 percent of hiPSC-CMs cultured under all conditions stained positive for ventricular myosin light chain (MLC-2V) (**Fig. S2B**).

**Figure 1.**
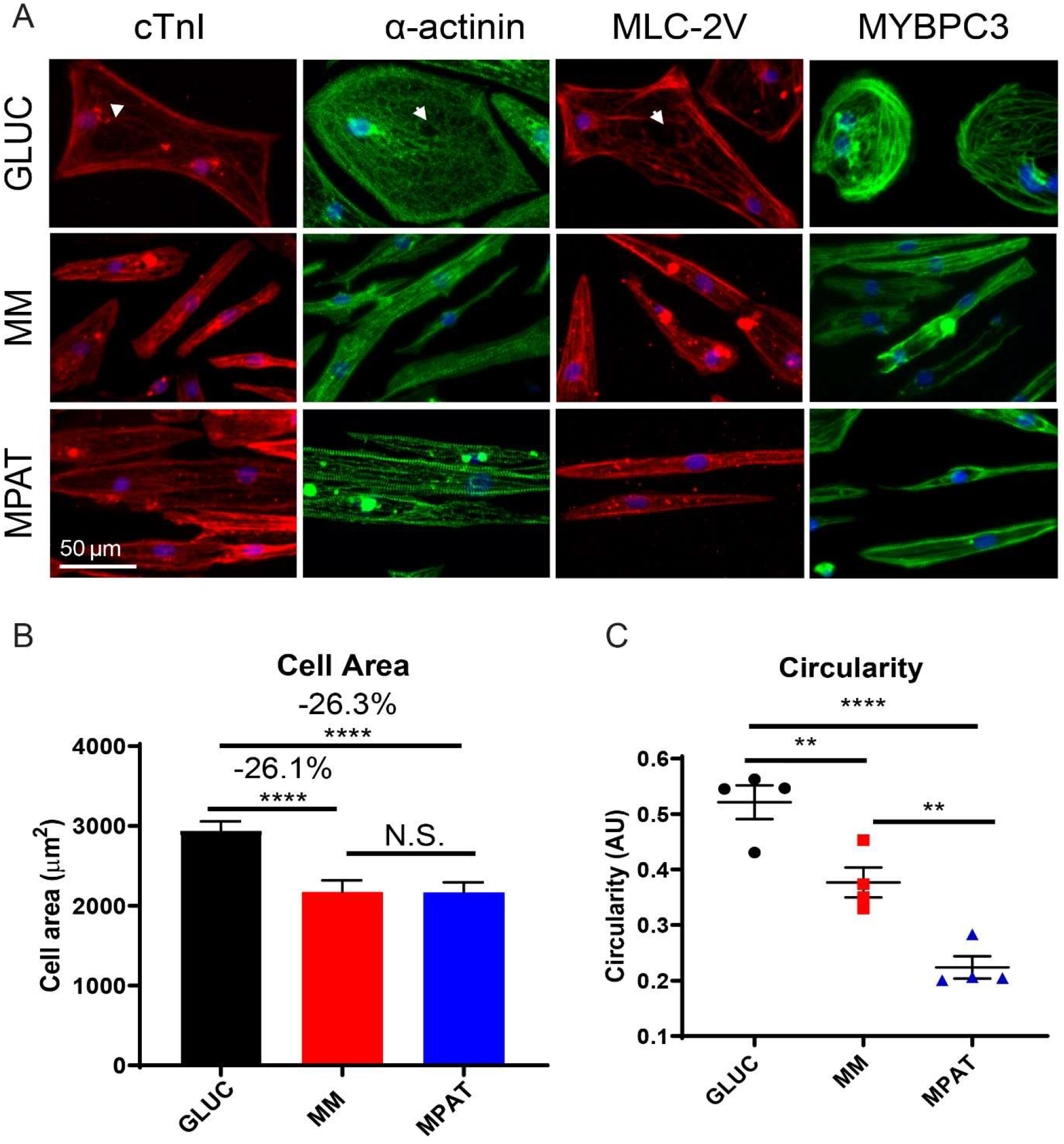
Morphological improvements with maturation methods in hiPSC-CMs. **A**, Representative images of staining for various sarcomeric proteins in hiPSC-CMs cultured in each condition. cTNI: Cardiac troponin I; α-actinin; MLC-2V: Myosin light chain, ventricular isoform; MYBPC3: myosin binding protein C3. White arrows indicate areas lacking sarcomeres. **B**, Average cell areas in hiPSC-CMs cultured under each condition. Data pooled from 3 independent experiments, 100-250 cells assessed per condition per experiment. **C**, Cellular circularity (defined by circularity × (4*π*area/perimeter^2^) in hiPSC-CMs cultured in each condition. Data from 4 independent experiments, 100-250 cells assessed per condition per experiment. **, ****, P<0.01, 0.0001, Kruskal-Wallis test with Dunn’s multiple comparison test (B), or one way ANOVA with Tukey’s multiple comparison test (C).

Using electron microscopy, we next examined sarcomeric morphology of hiPSC-CMs. When cultured in standard glucose medium, hiPSC-CMs displayed chaotically aligned, disorganized sarcomeres, as well as Z-lines of varying thickness, which often did not pass through the entirety of sarcomeres, similar to what we observed using immunofluorescence (**Fig 2. A-B**), and consistent with developing cardiomyocytes in the fetal heart^10^. By contrast, MM cells displayed more diverse sarcomere morphology: some sarcomeres were disordered, as seen in GLUC cells, whereas others were highly regular and organized, with I-bands, but lacking H-zones (**Fig. 2A-B**). MPAT cells consistently displayed organized sarcomeres in which both I-bands and H-zones could be observed (**Fig. 2A-B**), as is typically seen in more developed cardiomyocytes *in vivo*, or in more mature hiPSC-CMs^10, 11^. As the heart develops, a greater proportion of cardiomyocyte volume becomes occupied by myofibrils^12^, while average sarcomere length increases from approximately 1.6 to 2.2μm^2^. To assess whether changes in sarcomere occupancy were occurring in our cells, we measured mean α-actinin fluorescence intensity, and observed a significant (~45%) increase in α-actinin fluorescence in MPAT cells compared to GLUC (**Fig. S2C**). We also found that sarcomere length increased significantly, from ~1.81 to 1.92μm, from GLUC to MPAT (**Fig. 2C**). No significant increases in either α-actinin intensity or sarcomere length between MM and GLUC conditions were observed, suggesting that the MPAT condition induced cell maturity to the greatest degree.

**Figure 2.**
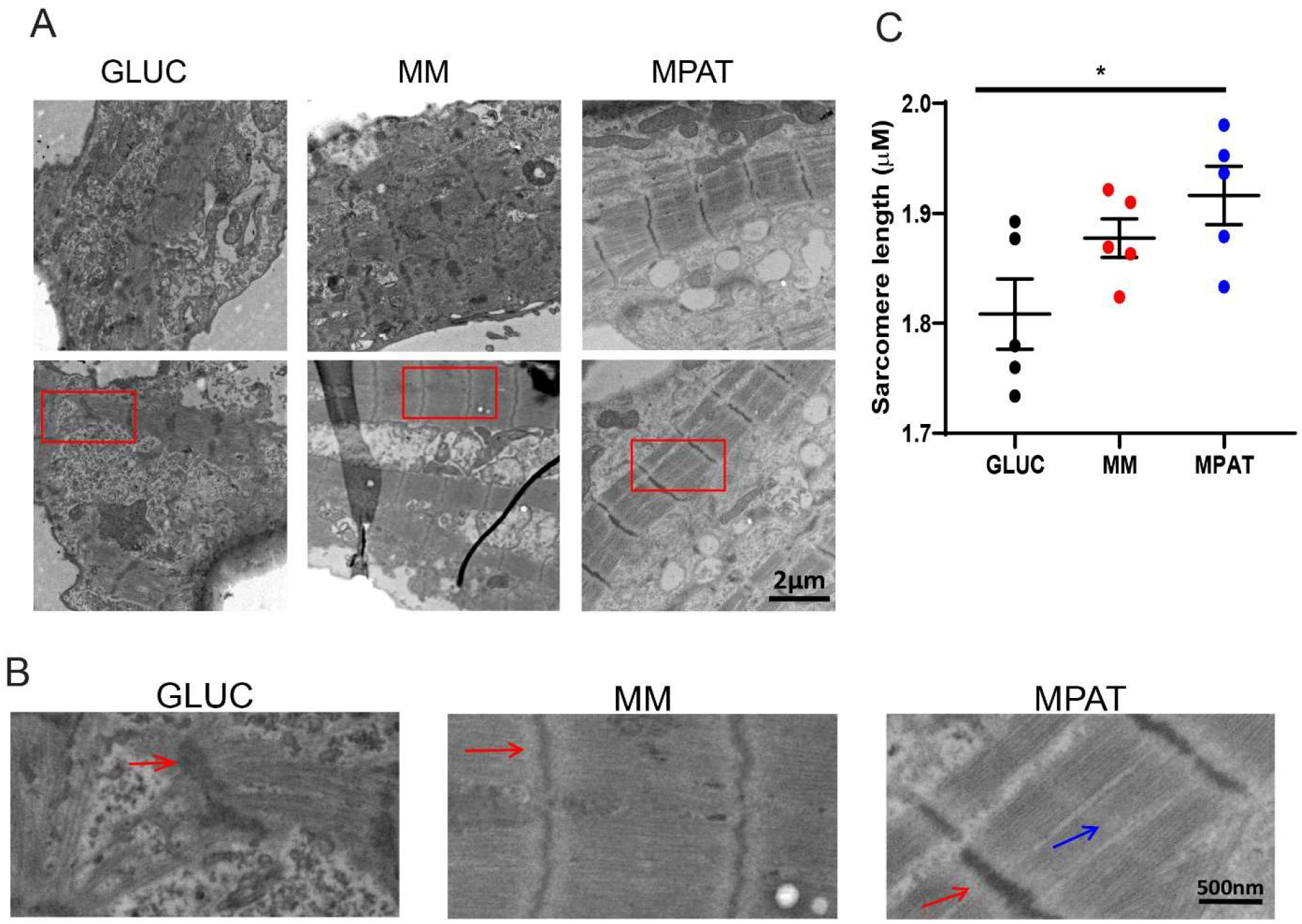
Sarcomere morphology is improved with maturity methods. **A-B**, Transmission election microscopy images of hiPSC-CMs cultured under each condition. Red boxes in (A) are enlarged in (B). Red arrows indicate I-bands, blue arrows indicate H-zones. **C**, Sarcomere length in hiPSC-CMs cultured in each condition. Sarcomeres measured between Z-lines, from 150-200 cells per condition per experiment, from 5 independent experiments. *P<0.05, one way ANOVA with Tukey’s multiple comparison test.

### Maturation methods improve myofibril force generation

Since our methods improved hiPSC-CM sarcomeric maturity and morphology, we investigated whether sarcomeric function was correspondingly improved. In order to compare sarcomeric function of hiPSC-CMs to adult cardiac tissue, we isolated myofibrils from hiPSC-CMs cultured under each condition and as well as from left ventricular tissue from a donor heart from a 35-year-old-male, as the donor of the MF750 hiPSC line was aged 32. Qualitatively, striations, indicative of sarcomeres, could be observed in some MPAT myofibrils, although not nearly to the extent of adult donor myofibrils, yet were typically absent from most GLUC and MM hiPSC-CM myofibrils (**Fig. 3A**). Remarkably, relative to GLUC, myofibrils from MPAT hiPSC-CMs demonstrated a nearly 150% increase in maximum tension generation, approaching the level of force generation measured the myofibrils of our adult donor heart (**Fig. 3C** and **Table S1**). MM myofibrils displayed an intermediate level of tension generation. Resting tension in GLUC, but not MM or MPAT myofibrils, was significantly lower than the resting tension from myofibrils isolated from donor hearts. Relaxation of myofibrils is biphasic, with a slow, linear phase, and a fast, exponential phase^13^. Activation and reactivation kinetics, slow and fast phase relaxation kinetics, and slow phase relaxation time were similar between all hiPSC-CM-derived and donor heart-derived myofibrils. We also conducted unsupervised hierarchical clustering on myofibril mechanical parameters in hiPSC-CMs and human myofibrils, and found that MPAT myofibrils clustered most closely with human adult myofibrils, potentially indicating increased maturity (**Fig. 3B**). To explore the mechanism behind increased MPAT force generation, we examined expression of myosin heavy chain isoforms and other myofibril-related genes on the protein level. We found that our cells expressed predominantly MYH7 (**Fig. S3A**). Expression of most other sarcomeric proteins was quite similar across various culture conditions. However, while MM and GLUC cells expressed both ventricular and atrial forms of myosin light chain (MLC-2V and MLC-2A), MPAT cells expressed primarily MLC-2V (**Fig. 3D**). Very low levels of slow skeletal troponin I, which is typically expressed in the fetal heart^14^, were detected in our samples. Significant expression of myosin IIB, which is expressed in neonatal but not adult heart^15^, was also detected. Taken together, these data indicate that the MPAT condition induces hiPSC-CM maturity in myofibril mechanics and sarcomeric protein expression.

**Figure 3.**
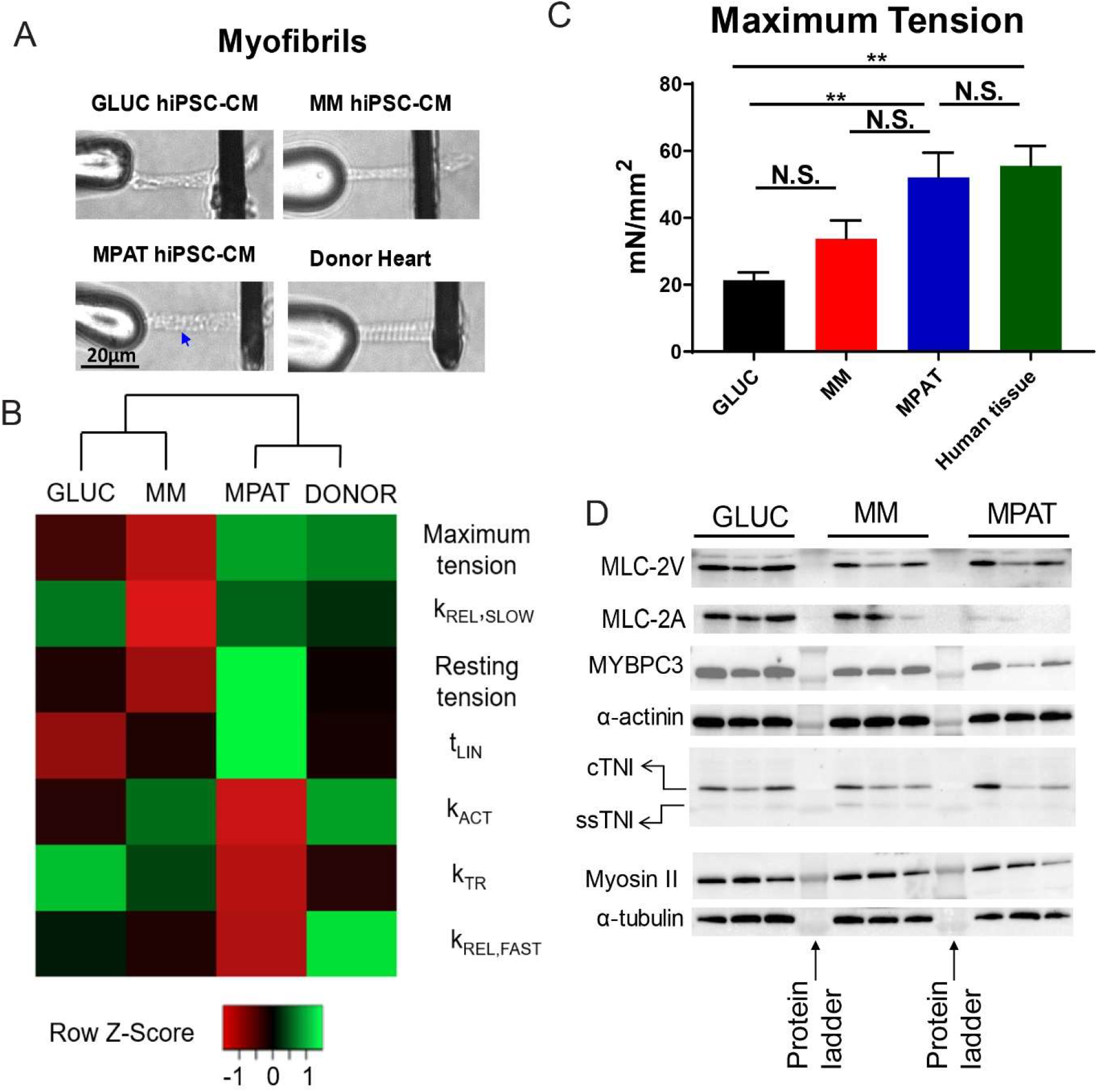
Myofibril mechanics in hiPSC-CMs and human heart tissue. **A**, Representative images of myofibrils isolated from each group and cells and from human donor hearts, mounted between stretcher (left) and force probe (right). Blue arrows indicates striations. **B**, Heatmap of hiPSC-CM myofibril mechanics. Unsupervised hierarchical clustering by centroid linkage performed on columns, indicating that MPAT cells cluster most closely with human adult donor myofibrils. **C**, Average myofibril maximum tension generation, normalized to myofibril cross sectional area. *,***, **** P<0.05, 0.001, 0.0001, one way ANOVA with Tukey’s multiple comparison test on log_2_-transformed data. 24-31 myofibrils were isolated from hiPSC-CMs, pooled from 4 independent cell inductions, while 6 myofibrils were isolated from the donor heart. For a complete set of myofibril mechanics data, see **Table S1**. **D**, Western blots on hiPSC-CMs lysates for myosin light chain ventricular isoform (MLC-2V), myosin light chain atrial isoform (MLC-2A), myosin binding protein C3 (MYBPC3), α-actinin, cardiac (C-TNI) and slow skeletal Troponin I (ss-TNI), myosin II, and α-tubulin as loading control. 4μg protein loaded per well.

### Maturation induces a gene expression program controlling fatty acid oxidation

To determine the mechanisms by which our methods induced cardiomyocyte maturation, RNA-seq was performed on hiPSC-CMs cultured under each condition. A large number of genes displayed differential expression between the GLUC and either MM or MPAT conditions, with a total of approximately 1000 genes showing either a 1.5-fold increase or decrease in expression (**Fig. S4A-B**). Most of the genes (733 upregulated and 529 downregulated genes) differentially regulated between either GLUC versus MM or GLUC versus MPAT were common between these two comparisons. Relatively few genes displayed differential expression between the MM and MPAT groups – only 234 genes were increased and 376 decreased by more than 1.5-fold (**Fig. S4A-B**). We also performed gene ontology (GO) analysis using PANTHER, and KEGG Pathway Analysis on differentially expressed genes. GO terms enriched in genes upregulated in MM or MPAT relative to GLUC hiPSC-CMs were frequently related to cardiac development or differentiation, or fatty acid metabolism, such as ‘regulation of heart morphogenesis’, ‘cardiocyte differentiation’, ‘fatty acid metabolism, and ‘fatty acid degradation’ (**Tables S2-3**). We also observed several terms related to cellular process development and elongation in the MM/MPAT upregulated genes, which could be related to the elongated cellular morphology we observe. Only a single GO term, ‘regulation of multicellular organismal process’ was enriched in MPAT relative to MM, while a few KEGG terms such as ‘adrenergic signaling in cardiomyocytes’ were enriched as well. By contrast, numerous GO and KEGG terms enriched in genes downregulated between MM versus GLUC or MPAT versus GLUC were related to DNA synthesis and cell division, including ‘cell cycle’, ‘cell division’, and ‘DNA replication’. Many of these terms were further downregulated in MPAT cells compared to MM. This could indicate that culture in fatty acid medium is arresting cell cycle entry and cell division in our hiPSC-CMs, which is consistent with a previous study in engineered heart tissue^16^.

Given that we observed differences in GO terms related to cardiac development or fatty acid metabolism, we investigated changes in relevant genes in our RNA-seq data. In fact, many genes involved in mitochondrial fatty acid uptake and long chain fatty acid oxidation (such as *CPT1A/1B/2* and *ACADVL*) were upregulated in MM, and further increased in MPAT hiPSC-CMs (**Fig. S4C**). To confirm our RNA-seq findings, we also assessed expression of these targets using qPCR, and found that several of these targets were significantly upregulated in MPAT cells, (**Fig. 4A**, with higher average levels of expression in MPAT cells than MM cells.

**Figure 4.**
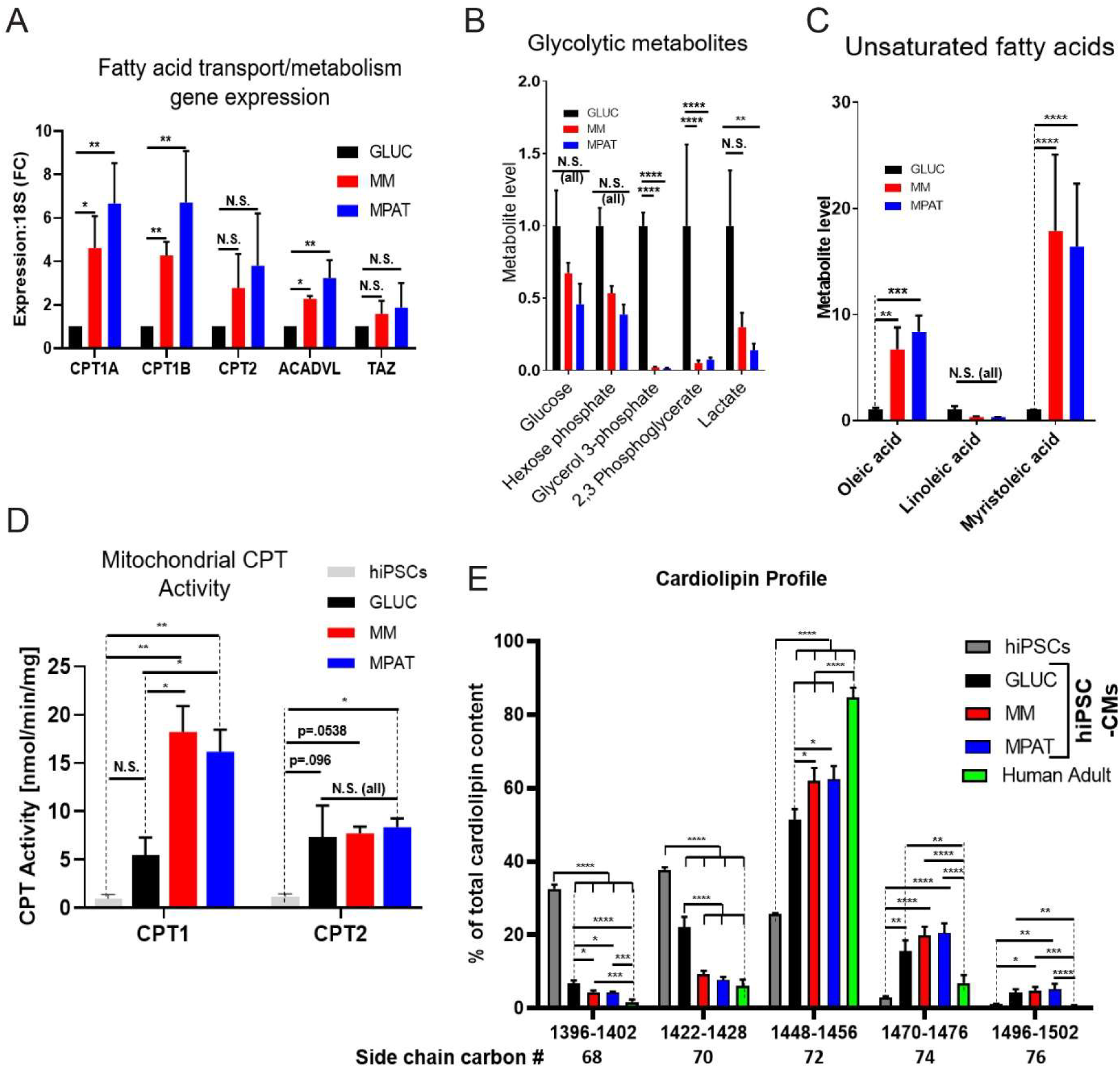
Maturation methods induce broad changes in hiPSC-CM metabolitic behavior. **A**, qPCR for selected genes involved in mitochondrial fatty acid uptake and metabolism and cardiolipin remodeling, normalized to 18S rRNA and shown as fold change relative to GLUC. **B-C**, Selected results from metabolic screening in hiPSC-CMS, parsed by metabolite type. Screening was performed on cells cultured in each maturation condition and expressed as fold change relative to GLUC hiPSC-CMs. Heatmaps of all assayed metabolites and of differentially expressed metabolites can be seen in **figure S5**. **D**, CPT1 and CPT2 activity in hiPSCs and hiPSC-CMs cultured under each condition. **E**, Quantification of peak areas of cardiolipin side chain percentages in hiPSCs and hiPSC-CMs cultured under each condition. *,**, ***, **** p<0.05. 0.01, 0.001, 0.0001, one way ANOVA followed by Tukey’s multiple comparison test. For **A-C**, tests performed on log_2_-transformed data. Cells or cDNA prepared from four independent inductions were used for **A-C**, while cells from three independent inductions and fifteen donor hearts were used for D and E.

To further assess metabolic changes, we performed metabolite screening on hiPSC-CMs cultured under each condition. We measured 136 polar metabolites in high throughput profiling, of which 51 were significantly altered across our three conditions. Levels of most metabolites were lower in MM and MPAT hiPSC-CMs relative to GLUC (**Fig. S5A-B**) while only 5 metabolites, methylmalonylcarnitine, O-tetradecanoyl-L-carnitine, tetradecenoyl-carnitine, catechol, and rhamnose, were significantly differentially produced when comparing MM and MPAT. We further examined metabolites on a candidate basis, focusing on glucose and long chain fatty acid metabolism. Surprisingly, glucose levels in hiPSC-CMs cultured under each condition were not significantly different (**Fig. 4B**), although glucose was lacking from maturation medium. However, many metabolites of the glycolytic pathway were significantly reduced in MPAT hiPSC-CMs relative to GLUC. Levels of most saturated fatty acids were similar among each group of hiPSC-CMs, including palmitate, an ingredient in MM. By contrast, intracellular levels of oleic acid, myristoleic acid, and several medium or long chain acylated fatty acids were much higher in MM and MPAT cells (**Fig. 4C, S5B**). Taken together with our RNA-seq data, these results indicate that MPAT induces a shift from glycolysis to fatty acid oxidation as a source of energy production, as is seen in the developing heart. Reduction in levels of glycolytic intermediates and products, and an increase in unsaturated fatty acids, may be driving the structural and functional maturation we observe in our cells.

### Maturation methods enhance mitochondrial CPT activity and cardiolipin maturation

In the developing heart, the switch from glycolysis to fatty acid oxidation is accompanied by an increase in activity of the carnitine palmitoyltransferase (CPT) system. CPT1A/B/C acylate fatty acids, allowing their import into the mitochondria, whereas CPT2 deacylates mitochondrial fatty acids to recover carnitine. As we observed increased expression of the CPT genes by RNA-seq and qPCR, we tested CPT activity in hiPSC-CMs and in undifferentiated hiPSCs. CPT1 activity was increased in all hiPSC-CM groups relative to hiPSCs, and was further increased in MM/MPAT groups (**Fig. 4D**). CPT2 activity was similar between all groups of hiPSC-CMs, but was significantly increased only in MPAT hiPSC-CMs relative to hiPSCs. As CPT1 activity is considered to be the rate limiting step in long chain fatty acid oxidation^17^, increased CPT1 activity is consistent with increased use of long chain fatty acids, and is likely indicative of metabolic maturation.

Cardiolipin (CL) is the critical phospholipid of mitochondrial membranes, and is essential for mitochondrial function^18^. Each CL molecule has four fatty acid side chains. Side chain composition varies by tissue type, with the majority of CL in the adult heart having unsaturated fatty acid side chains, with tetralineolyl CL (total molecular weight of 1448 Daltons) by far the most abundant species^19^. CL side chain composition is important for metabolic function in the heart, and defects in CL sidechain remodeling in the heart have been associated with Barth Syndrome^20^. In the developing heart, CL content changes, with an increase in CL having side chains with 72 total carbons (72C-CL, MW 1448-1456) and a loss of most other species. Considering the metabolic improvements we observed in our MPAT cells, we investigated whether CL remodeling was occurring in our more mature cells, using adult human hearts as control. Induction from hiPSCs into hiPSC-CMs caused significant CL remodeling, in particular a decrease in 68C- and 70C-CL, and an increase in 72C- and 74C-CL. With maturation methods, a further loss in 70C-CL and increases in both 72C- and 74C-CL were observed (**Figs. 4E** and **S6**). These changes shifted the CL profile towards that of adult human hearts. We also assessed expression of Tafazzin (Taz), a major enzyme involved in CL remodeling in muscle tissues^18^, but did not observe changes in its expression (**Fig. 4A**).

### Maturation methods allow an adult like hypertrophic response in hiPSC-CMs

We next investigated the efficacy of our maturation methods for studying hypertrophic remodeling. Significant differences in the magnitude of the *in vitro* hypertrophic response have been observed in neonatal versus adult cardiac myocytes. In neonatal cardiomyocytes, agents such as the alpha-adrenergic receptor agonist phenylephrine (PE) induce a 50-100% in increase in cell area^21, 22^, whereas the response in adult cardiomyocytes is typically only 10-30% ^21, 23, 24^. Therefore, we investigated the response of hiPSC-CMs to PE. Since the BET-bromodomain inhibitor JQ1 is known to inhibit cardiomyocyte hypertrophy both in vivo and *in vitro*^22, 25, 26^; we also tested whether JQ1 could block the effects of PE in our system. Surprisingly, PE treatment had no effect on cell area of GLUC hiPSC-CMs, yet JQ1 treatment nonetheless profoundly reduced basal cell area (**Fig. 5A-B**). However, in MM and MPAT hiPSC-CMs, PE treatment induced a strong hypertrophic response, causing a 29% increase in cell area in MM and a 19% increase in MPAT cells. In both groups, this hypertrophy was blocked by JQ1 treatment: cell area relative to PE-treated cells was reduced by 31% in MPAT and 17% in MM cells. To validate these findings, we also investigated expression of *NPPB* (BNP), a robust marker of hiPSC-CM hypertrophy^27^ by qPCR and immunofluorescence. By qPCR, MM and MPAT cells displayed an approximately 5-6 fold reduction in BNP expression relative to GLUC cells (**Fig. 5C**). Furthermore, more than 20% of GLUC cells stained positive for BNP expression, which was not increased by PE treatment. Basal BNP staining in MM and MPAT cells was much lower, yet was increased 2 to 2.5 fold by PE treatment (**Fig. S7A-B**). Taken together, these results suggest that prolonged glucose culture may be inducing hiPSC-CM hypertrophy, and that proper selection of culture conditions may be essential to characterizing hypertrophic cardiomyopathy using the hiPSC-CM platform.

**Figure 5.**
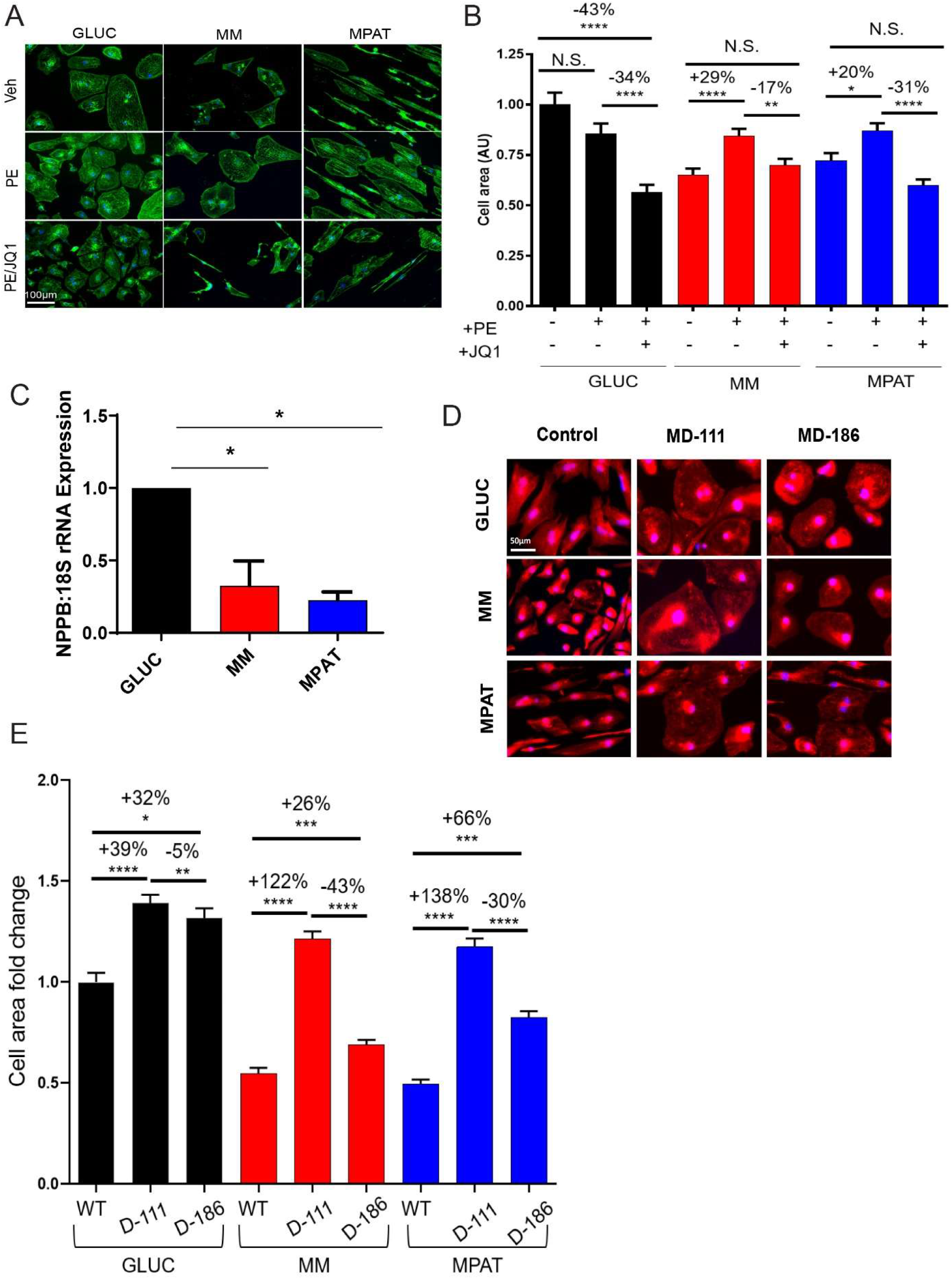
Culture in maturation medium allows induction of a hypertrophic response in hiPSC-CMs. **A-B**, hiPSC-CMs cultured under indication conditions and treated with vehicle, 10 μmol/L phenylephrine (PE) or 10 μmol/L PE and 1 μmol/L JQ1 (a BET Bromodomain inhibitor) for 48 hours, then fixed and stained for α-actinin (green) and Hoechst 33342 (blue). Representative images are shown in **A**. Cell area of hiPSC-CMs were quantified in **B**. Cell areas measured on 616-789 cardiomyocytes pooled from four independent inductions. **C**, qPCR for B-type natriuretic peptide, normalized to 18S rRNA and shown as fold change relative to GLUC, from cDNA prepared from four independent inductions. **D-F**, Characterization of cardiac hypertrophy in hiPSC-CMs derived from either a control line or from patients with Danon disease (MD-111, MD-186), and cultured under the indicated conditions. **D**, Representative images of hiPSC-CMs stained with CellMask Orange (red) and Hoechst 33342 (blue). **E**, Quantification of cell area, measured on 356-1056 cardiomyocytes pooled from two independent inductions. *, **, **** p<0.05, 0.01, 0.0001, Kruskal-Wallis test followed by Dunn’s multiple comparison test (B,E) or one way ANOVA followed by Tukey’s multiple comparison test on log_2_ transformed data (C).

To compare our hiPSC-CMs directly to adult cells, we also isolated adult mouse ventricular cardiomyocytes (AMVCMS), and treated them with PE and JQ1. In AMVCMS, PE induced a 16% increase in cell area, while JQ1 induced an 11% decrease (**Fig. S7C-D**).

We next investigated whether we could characterize cardiac hypertrophy in patient-specific cells. In a previous study^28^, we generated multiple lines of hiPSC-CMs from Danon disease patients carrying mutations in the *LAMP2* gene, leading to pathological cardiac remodeling. Interestingly, echocardiography indicated that the patients from whom we generated hiPSC-CMs displayed varying degrees of cardiac hypertrophy and dysfunction. We therefore focused on hiPSC-CMs from two patients, MD-186 and MD-111, with moderate and severe cardiac hypertrophy, respectively. The left ventricular posterior wall thickness (LVPW) of patient MD-111 was 18mm, whereas the LVPW of MD-186 was 10mm^28^. Compared to previously reported LVPW values in healthy adult males (6-10mm, mean 8mm),^29^ these values represent a 125% and 20% increase, respectively. We found that when cultured in GLUC media, both sets of Danon hiPSC-CM lines displayed a modest and similar increase in cell area (30-40% increase), compared to hiPSC-CMs derived from the MF750 control line (**Fig. 5D-E**). However, when cultured in MPAT or MM, the two Danon cell lines displayed differential degrees of hypertrophy: MD-111 hiPSC-CMs displayed extreme hypertrophy (138% increase in MPAT), whereas the hypertrophy of MD-186 cells was comparatively modest (66% increase in MPAT). The extent of hypertrophic remodeling observed in these two cell lines under maturation methods correlated relatively well to the increase in left ventricular posterior wall thickness (LVPW) observed in the hearts of patients MD-186 and MD-111, suggesting the possibility of recapitulating patient-specific phenotypes using more mature cells.

### PE treatment of mature hiPSC-CMs induces myofibril relaxation changes as observed in hypertrophic cardiomyopathy

Previous reports have indicated that myofibrils isolated from hearts of humans with hypertrophic cardiomyopathy (HCM) display profound differences in mechanical behavior^30^. We examined whether myofibrils isolated from hypertrophic hiPSC-CMs might demonstrate similar myofibril mechanical perturbations. For comparison, we first assessed myofibril mechanics from human donor and HCM hearts in our tissue bank. Myofibrils were isolated from 4 donor hearts and 4 hearts from patients transplanted for hypertrophic cardiomyopathy (HCM) with a left ventricular posterior wall thickness of >12mm as assessed by echocardiography. All myofibril mechanical parameters from donor and HCM hearts are listed in **Table S5**. Compared to donor hearts, HCM hearts-derived myofibrils showed a shortened linear phase relaxation time, as well as faster activation kinetics (**Fig. 6A**). These results are similar to findings from myofibrils isolated from a human heart with an HCM-associated mutation in MYH7^30^.

**Figure 6:**
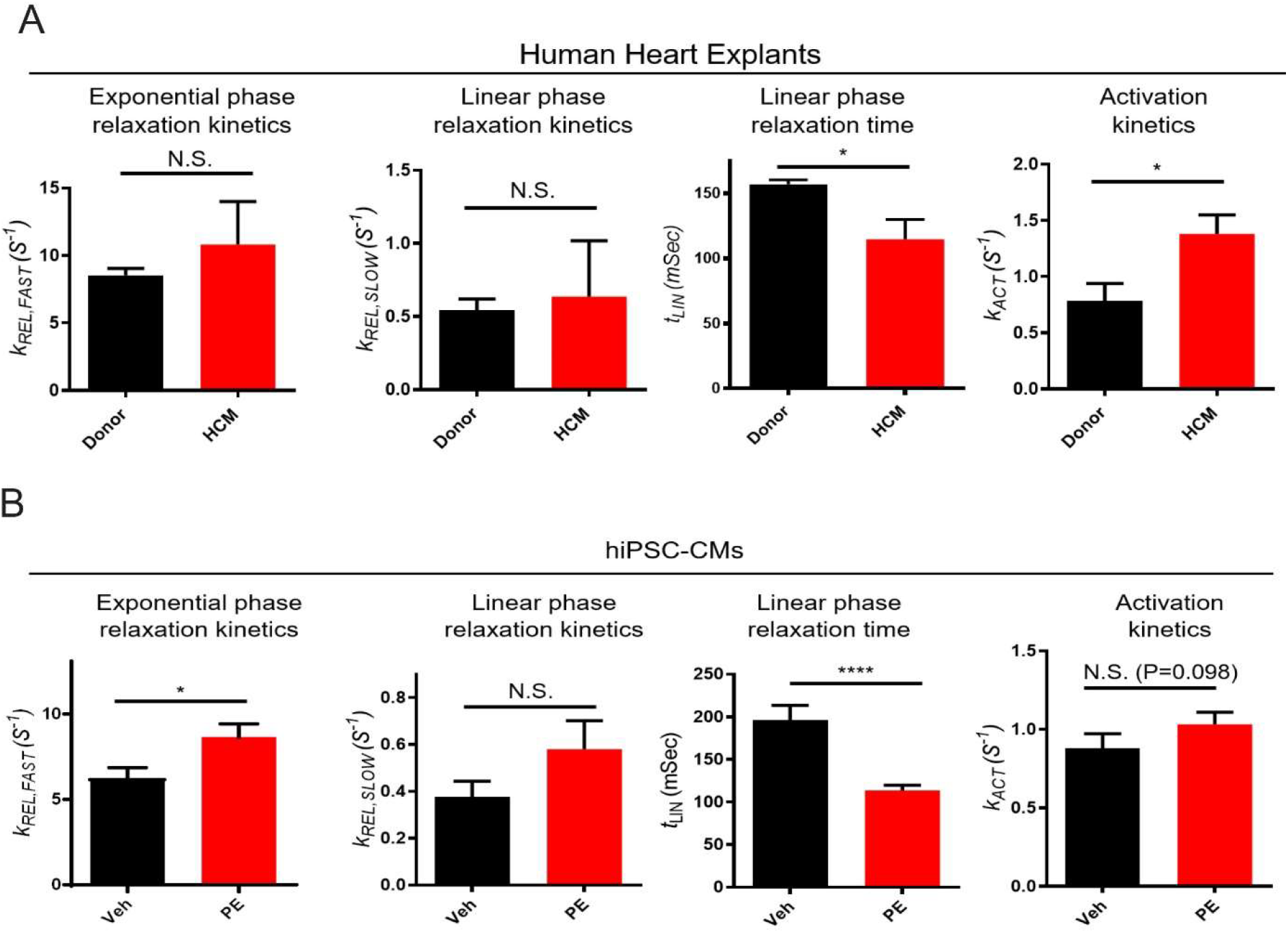
Phenylephrine treatment induces changes in myofibril relaxation which are similar to those observed in human hypertrophic cardiomyopathy. **A**, Myofibril mechanics data from human donor and hypertrophic cardiomyopathy (HCM) hearts. Myofibrils were isolated from 4 donor and 4 HCM hearts, with at least 6 myofibrils isolated per heart. **B**, MPAT hiPSC-CMs were treated with either vehicle or 10 μmol/L PE for 48 hours. Myofibril mechanics data from MPAT hiPSC-CMs treated with either vehicle or 10 μmol/L PE for 48 hours. 44-45 myofibrils isolated per condition, pooled from 3 independent inductions. Complete tables of myofibril data from these experiments are available in **Tables S4 and S5**. *, **, **** p<0.05, 0.01, 0.0001, Student’s t test on log_2_-transformed data.

Myofibrils were then isolated from hypertrophic hiPSC-CMs. For this experiment, we chose to isolate myofibrils from MPAT hiPSC-CMs, as these displayed the most adult-like myofibril behavior. Hypertrophy was again induced via treatment with PE. To compare hiPSC-CMs to hypertrophic adult cardiomyocytes, we also isolated myofibrils from vehicle or PE-treated AMVCMs. Remarkably, we found that myofibril relaxation was altered in the PE-treated hiPSC-CMs – the exponential phase relaxation constant was increased, while the duration of linear phase relaxation was shortened (indicating faster relaxation) (**Fig. 6B**). Activation kinetics also trended towards an increase. In AMVCMs, PE induced similar relaxation changes, but no changes in activation kinetics could be detected (**Fig. S9**). The changes in myofibril relaxation induced by PE treatment were relatively similar to those observed in human HCM. Together, these results indicate that the MPAT hiPSC-CM platform is a suitable *in vitro* model for human hypertrophic cardiomyopathy, allowing parallel assessment of changes in cell area, hypertrophic gene expression, and myofibril mechanical behavior.

## Discussion

Here, we present a relatively simple method to produce more mature hiPSC-CMs by culturing cells in medium containing fatty acids as a significant energy source and plating these cells on micropatterned growth surfaces. Under these conditions, cells demonstrate elongation, enhanced sarcomeric maturity, metabolic gene expression, mitochondrial fatty acid uptake, and cardiolipin maturation. The mechanical behavior of myofibrils isolated from these cells more closely resembles those from adult human ventricular tissue, particularly in terms of force generation. These mature hiPSC-CMs also respond well to hypertrophic stimulation, producing an adult-like hypertrophic response, and demonstrating alterations in myofibrils relaxation similar to human HCM tissue.

Improving hiPSC-CM maturity has recently been a topic of acute interest, with dozens of studies reporting a plethora of methods of achieving this aim, including prolonged culture, electrical pacing, treatment with T3 hormone, and engineered heart tissue setups^2^. Compared to many of these approaches, our combinatorial approach may have potential applications due to its simplicity. Fatty acid medium can be prepared at low cost, while patterned surfaces require only a few simple tools to produce. This culturing setup can therefore be introduced into a new research environment quite quickly. The combination of these methods induces a more mature cardiac phenotype across a variety of parameters, from cell morphology (**Figs. 1, 2**) to contractility (**Figs. 3, 6**), and hypertrophic response (**Figs. 5, 6, S7**). It is interesting to note that the two maturation steps used seem to exert their benefits on different aspects of cardiac maturity. For example, most of the improvements in myofibril mechanics occurred in MPAT compared to MM hiPSC-CMs. This indicates that combinatorial approaches may be necessary to produce mature hiPSC-CMs. We believe that the method we detailed here has a number of exciting future applications, from screening for new cardiac drugs to interrogating the effect of genetic or pharmacological perturbations in a human cell system. The approaches used here could also be combined with methods such as cell pacing, growth hormone treatment, or use of altered ECM composition or stiffness to promote further maturation.

While our methods appear to drive hiPSC-CMs towards maturity, the mechanisms by which this occurs have not been fully elucidated. Metabolic screening data indicated that the most altered metabolites between conditions were intermediates and products of glycolysis (higher in less mature cells), oleic and myristoleic acid, and intermediates of long chain fatty acid oxidation (higher in more mature cells) (**Fig. 4**). This may indicate that a shift from glycolytic energy production to a reliance on fatty acids is a critical part of this process. We also observed changes in cardiolipin content, but whether this is a cause or consequence of greater maturity remains unknown. The maturity-inducing effects of micro and nanopatterned substrates on hiPSC-CMs are well reported^3, 31^, but the mechanism driving this remains undefined. A previous report indicated that a 700-1000nm nanopatterned groove size is optimal to induce hiPSC-CM maturity^31^. While the micropatterned grooves described here are much larger, they clearly exert positive effects on maturity as well. Our results indicate that patterned surfaces primarily alter cellular morphology (**Fig. 1C**), sarcomere organization (**Fig. 2**), and myofibril behavior (**Fig. 3**), but we also observed a degree of CL remodeling and increased fatty acid gene expression in MPAT versus MM cells, (**Fig. 4**). Immature hiPSC-CMs and fetal cardiomyocytes typically have an atrial cell-like identity, with dominant expression of the atrial isoforms of myosin light and heavy chains, MLC-2A and MYH6, respectively^2, 32^. We observed predominant expression of MYH7 in all hiPSC-CM cultures, and increased relative expression of MLC-2V to MLC-2A in MPAT hiPSC-CMs. The activation kinetics we measured in hiPSC-CMs myofibrils (0.95-1.04) are lower than previously reported values in human atrial tissue myofibrils (3.73)^33^ or in eSC-CM myofibrils expressing either mostly MYH7 or mixed MYH6/7 (1.70 or 2.44 respectively).^34^ Together, these findings are indicative of sarcomeric maturity and potentially a more ventricular-like identity in our MPAT cells. As forced expression of MLC-2V results in increased contractility in isolated cardiomyocytes^35^, this could be a mechanism underlying the increased myofibril force generation we observe in MPAT cells.

Previous reports have indicated that prolonged (> 100 days) culture of hiPSC-CMs induces a more adult-like phenotype, as indicated by improvements in sarcomeric ultrastructure, cell morphology, and contractility^11, 36^. Yet, hiPSC-CMs cultured in glucose-containing media had no hypertrophic response to PE, high levels of BNP expression, and modest induction of hypertrophy in Danon hiPSC-CMs (**Fig. 5, S7**). The antihypertrophic agent JQ1 also reduced cell area significantly beyond baseline in GLUC hiPSC-CMs, which did not occur in MM or MPAT cells (**Fig. 5A-B**). A recent report indicated that 14 days of culture in 22 mM glucose was sufficient to induce cardiac hypertrophy and dysfunction in hiPSC-CMs^37^. Although our GLUC medium contains only 10.9 mM glucose, prolonged culture under these conditions may be sufficient to induce a hypertrophic phenotype. Similarly, it has been reported that supplementation of culture medium with either 5 or 20% fetal calf serum for one week masked the prohypertrophic effects of PE treatment in hiPSC-CMs 30-40 days post induction^38^. Myofibrils isolated from GLUC cells also demonstrated low levels of active tension generation. This could indicate lack of maturity, but could also be a product of their hypertrophic phenotype: for example, certain forms of cardiac dysfunction, such as Danon disease, are associated with reduced active tension generation^28^. hiPSC-CMs are increasingly used to model various types of cardiomyopathies, and these studies, along with our findings, indicate that a careful choice of conditions for long term culture is essential for *in vitro* characterization of hypertrophic cardiomyopathy.

While we observed similar changes in relaxation in PE-treated hiPSC-CMs and human HCM hearts, PE treatment did not dramatically induce an increase of activation kinetics. This could be due to differences between the myofibrils from human heart and hiPSC-CMs, as hiPSC-CMs show higher baseline activation kinetics (**Table S1** and **S4** versus **S5**). However, 48 hours of PE treatment is also unlikely to fully recapitulate the remodeling which occurs in HCM, which is mutation driven and occur over years. Studying myofibril mechanics in hiPSC-CMs with HCM-causing mutations could therefore be of interest for future studies. A recent report demonstrated that 48 hours of PE treatment on adult rat cardiac (ARVM) myofibrils did not affect the kinetics of myofibril activation or relaxation^39^. Here, we demonstrated that 72 hours of PE treatment on AMVCMs resulted in a slight (~10ms) reduction in linear phase relaxation time. PE treatment had greater effects in hiPSC-CMs, where linear phase was shortened by more than 80 ms (**Fig. S8, Table S4, S6**). It is possible that these divergent effects are due to differences between the PE response of mouse, rat, and human, or differences in treatment times. However, the kinetics of human and rodent myofibrils are also dissimilar: in rodent, activation and relaxation constants are much larger, and linear phase relaxation time is much shorter (~50 ms in rodents versus 150-200 ms in human). The capability of PE to further speed up this already short linear phase may therefore be abrogated, leading to modest effects in rodent.

In totality, our findings suggest that matured hiPSC-CMs are an appropriate *in vitro* model to conduct translational research of human myofibril mechanics in response to pathological stresses. The platform we describe here could also be of particular use when combined with CRISPR-based mutation strategies to create cardiomyopathy patient-specific cells, allowing simultaneous assessment of cellular morphology, myofibril mechanics, metabolic behavior, and hypertrophic remodeling.

## Abbreviations

CL: Cardiolipin
CPT: Carnitine palmitoyltransferase
GLUC: Glucose cells: hiPSC-CMs cultured in glucose-based media
hiPSC-CMs: Human pluripotent stem cell-derived cardiomyocytes
MM: Maturation medium: hiPSC-CMs cultured in fatty acid-based media
MPAT: Maturation medium/patterned cells: hiPSC-CMs cultured on patterned surfaces in fatty acid-based media
MYH: myosin heavy chain
MYL/MLC: myosin light chain

## Supplemental Materials

Expanded Materials & Methods

Online Figures I-IX

Online Tables S1-S6

Online Video I-III Legends

References 40-53

## References

1. Mills RJ and Hudson JE. Bioengineering adult human heart tissue: How close are we? APL Bioeng. 2019;3:010901.

2. Tan SH and Ye L. Maturation of Pluripotent Stem Cell-Derived Cardiomyocytes: a Critical Step for Drug Development and Cell Therapy. J Cardiovasc Transl Res. 2018;11:375–392.

3. Pioner JM, Racca AW, Klaiman JM, Yang KC, Guan X, Pabon L, Muskheli V, Zaunbrecher R, Macadangdang J, Jeong MY, Mack DL, Childers MK, Kim DH, Tesi C, Poggesi C, Murry CE and Regnier M. Isolation and Mechanical Measurements of Myofibrils from Human Induced Pluripotent Stem Cell-Derived Cardiomyocytes. Stem Cell Reports. 2016;6:885–96.

4. Poon E, Keung W, Liang Y, Ramalingam R, Yan B, Zhang S, Chopra A, Moore J, Herren A, Lieu DK, Wong HS, Weng Z, Wong OT, Lam YW, Tomaselli GF, Chen C, Boheler KR and Li RA. Proteomic Analysis of Human Pluripotent Stem Cell-Derived, Fetal, and Adult Ventricular Cardiomyocytes Reveals Pathways Crucial for Cardiac Metabolism and Maturation. Circ Cardiovasc Genet. 2015;8:427–36.

5. Lian X, Hsiao C, Wilson G, Zhu K, Hazeltine LB, Azarin SM, Raval KK, Zhang J, Kamp TJ and Palecek SP. Robust cardiomyocyte differentiation from human pluripotent stem cells via temporal modulation of canonical Wnt signaling. Proc Natl Acad Sci U S A. 2012;109:E1848–57.

6. Tohyama S, Hattori F, Sano M, Hishiki T, Nagahata Y, Matsuura T, Hashimoto H, Suzuki T, Yamashita H, Satoh Y, Egashira T, Seki T, Muraoka N, Yamakawa H, Ohgino Y, Tanaka T, Yoichi M, Yuasa S, Murata M, Suematsu M and Fukuda K. Distinct metabolic flow enables large-scale purification of mouse and human pluripotent stem cell-derived cardiomyocytes. Cell Stem Cell. 2013;12:127–37.

7. Correia C, Koshkin A, Duarte P, Hu D, Teixeira A, Domian I, Serra M and Alves PM. Distinct carbon sources affect structural and functional maturation of cardiomyocytes derived from human pluripotent stem cells. Sci Rep. 2017;7:8590.

8. Hu D, Linders A, Yamak A, Correia C, Kijlstra JD, Garakani A, Xiao L, Milan DJ, van der Meer P, Serra M, Alves PM and Domian IJ. Metabolic Maturation of Human Pluripotent Stem Cell-Derived Cardiomyocytes by Inhibition of HIF1alpha and LDHA. Circ Res. 2018;123:1066–1079.

9. Brewer GJ and Cotman CW. Survival and growth of hippocampal neurons in defined medium at low density: advantages of a sandwich culture technique or low oxygen. Brain Res. 1989;494:65–74.

10. Zhang F and Pasumarthi KB. Ultrastructural and immunocharacterization of undifferentiated myocardial cells in the developing mouse heart. J Cell Mol Med. 2007;11:552–60.

11. Lundy SD, Zhu WZ, Regnier M and Laflamme MA. Structural and functional maturation of cardiomyocytes derived from human pluripotent stem cells. Stem Cells Dev. 2013;22:1991–2002.

12. Olivetti G, Anversa P and Loud AV. Morphometric study of early postnatal development in the left and right ventricular myocardium of the rat. II. Tissue composition, capillary growth, and sarcoplasmic alterations. Circ Res. 1980;46:503–12.

13. Stehle R, Kruger M and Pfitzer G. Force kinetics and individual sarcomere dynamics in cardiac myofibrils after rapid ca(2+) changes. Biophys J. 2002;83:2152–61.

14. Saggin L, Gorza L, Ausoni S and Schiaffino S. Troponin I switching in the developing heart. J Biol Chem. 1989;264:16299–302.

15. Ma X and Adelstein RS. In vivo studies on nonmuscle myosin II expression and function in heart development. Front Biosci (Landmark Ed). 2012;17:545–55.

16. Mills RJ, Titmarsh DM, Koenig X, Parker BL, Ryall JG, Quaife-Ryan GA, Voges HK, Hodson MP, Ferguson C, Drowley L, Plowright AT, Needham EJ, Wang QD, Gregorevic P, Xin M, Thomas WG, Parton RG, Nielsen LK, Launikonis BS, James DE, Elliott DA, Porrello ER and Hudson JE. Functional screening in human cardiac organoids reveals a metabolic mechanism for cardiomyocyte cell cycle arrest. Proc Natl Acad Sci U S A. 2017;114:E8372–E8381.

17. Noland RC. Exercise and Regulation of Lipid Metabolism. Prog Mol Biol Transl Sci. 2015;135:39–74.

18. Shen Z, Ye C, McCain K and Greenberg ML. The Role of Cardiolipin in Cardiovascular Health. Biomed Res Int. 2015;2015:891707.

19. Schlame M, Ren M, Xu Y, Greenberg ML and Haller I. Molecular symmetry in mitochondrial cardiolipins. Chem Phys Lipids. 2005;138:38–49.

20. Schlame M, Towbin JA, Heerdt PM, Jehle R, DiMauro S and Blanck TJ. Deficiency of tetralinoleoyl-cardiolipin in Barth syndrome. Ann Neurol. 2002;51:634–7.

21. Miller CL, Oikawa M, Cai Y, Wojtovich AP, Nagel DJ, Xu X, Xu H, Florio V, Rybalkin SD, Beavo JA, Chen YF, Li JD, Blaxall BC, Abe J and Yan C. Role of Ca2+/calmodulin-stimulated cyclic nucleotide phosphodiesterase 1 in mediating cardiomyocyte hypertrophy. Circ Res. 2009;105:956–64.

22. Anand P, Brown JD, Lin CY, Qi J, Zhang R, Artero PC, Alaiti MA, Bullard J, Alazem K, Margulies KB, Cappola TP, Lemieux M, Plutzky J, Bradner JE and Haldar SM. BET bromodomains mediate transcriptional pause release in heart failure. Cell. 2013;154:569–82.

23. Knight WE, Chen S, Zhang Y, Oikawa M, Wu M, Zhou Q, Miller CL, Cai Y, Mickelsen DM, Moravec C, Small EM, Abe J and Yan C. PDE1C deficiency antagonizes pathological cardiac remodeling and dysfunction. Proc Natl Acad Sci U S A. 2016.

24. Bupha-Intr T, Haizlip KM and Janssen PM. Role of endothelin in the induction of cardiac hypertrophy *in vitro*. PLoS One. 2012;7:e43179.

25. Duan Q, McMahon S, Anand P, Shah H, Thomas S, Salunga HT, Huang Y, Zhang R, Sahadevan A, Lemieux ME, Brown JD, Srivastava D, Bradner JE, McKinsey TA and Haldar SM. BET bromodomain inhibition suppresses innate inflammatory and profibrotic transcriptional networks in heart failure. Sci Transl Med. 2017;9.

26. Spiltoir JI, Stratton MS, Cavasin MA, Demos-Davies K, Reid BG, Qi J, Bradner JE and McKinsey TA. BET acetyl-lysine binding proteins control pathological cardiac hypertrophy. J Mol Cell Cardiol. 2013;63:175–9.

27. Carlson C, Koonce C, Aoyama N, Einhorn S, Fiene S, Thompson A, Swanson B, Anson B and Kattman S. Phenotypic screening with human iPS cell-derived cardiomyocytes: HTS-compatible assays for interrogating cardiac hypertrophy. J Biomol Screen. 2013;18:1203–11.

28. Chi C, Leonard A, Knight WE, Beussman KM, Zhao Y, Cao Y, Londono P, Aune E, Trembley MA, Small EM, Jeong MY, Walker LA, Xu H, Sniadecki NJ, Taylor MR, Buttrick PM and Song K. LAMP-2B regulates human cardiomyocyte function by mediating autophagosome-lysosome fusion. Proc Natl Acad Sci U S A. 2018.

29. Lang RM, Badano LP, Mor-Avi V, Afilalo J, Armstrong A, Ernande L, Flachskampf FA, Foster E, Goldstein SA, Kuznetsova T, Lancellotti P, Muraru D, Picard MH, Rietzschel ER, Rudski L, Spencer KT, Tsang W and Voigt JU. Recommendations for cardiac chamber quantification by echocardiography in adults: an update from the American Society of Echocardiography and the European Association of Cardiovascular Imaging. J Am Soc Echocardiogr. 2015;28:1–39 e14.

30. Belus A, Piroddi N, Scellini B, Tesi C, D’Amati G, Girolami F, Yacoub M, Cecchi F, Olivotto I and Poggesi C. The familial hypertrophic cardiomyopathy-associated myosin mutation R403Q accelerates tension generation and relaxation of human cardiac myofibrils. J Physiol. 2008;586:3639–44.

31. Carson D, Hnilova M, Yang X, Nemeth CL, Tsui JH, Smith AS, Jiao A, Regnier M, Murry CE, Tamerler C and Kim DH. Nanotopography-Induced Structural Anisotropy and Sarcomere Development in Human Cardiomyocytes Derived from Induced Pluripotent Stem Cells. ACS Appl Mater Interfaces. 2016;8:21923–32.

32. Bedada FB, Wheelwright M and Metzger JM. Maturation status of sarcomere structure and function in human iPSC-derived cardiac myocytes. Biochim Biophys Acta. 2016;1863:1829–38.

33. Piroddi N, Belus A, Scellini B, Tesi C, Giunti G, Cerbai E, Mugelli A and Poggesi C. Tension generation and relaxation in single myofibrils from human atrial and ventricular myocardium. Pflugers Arch. 2007;454:63–73.

34. Weber N, Schwanke K, Greten S, Wendland M, Iorga B, Fischer M, Geers-Knorr C, Hegermann J, Wrede C, Fiedler J, Kempf H, Franke A, Piep B, Pfanne A, Thum T, Martin U, Brenner B, Zweigerdt R and Kraft T. Stiff matrix induces switch to pure beta-cardiac myosin heavy chain expression in human ESC-derived cardiomyocytes. Basic Res Cardiol. 2016;111:68.

35. Pawloski-Dahm CM, Song G, Kirkpatrick DL, Palermo J, Gulick J, Dorn GW, 2nd, Robbins J and Walsh RA. Effects of total replacement of atrial myosin light chain-2 with the ventricular isoform in atrial myocytes of transgenic mice. Circulation. 1998;97:1508–13.

36. Kamakura T, Makiyama T, Sasaki K, Yoshida Y, Wuriyanghai Y, Chen J, Hattori T, Ohno S, Kita T, Horie M, Yamanaka S and Kimura T. Ultrastructural maturation of human-induced pluripotent stem cell-derived cardiomyocytes in a long-term culture. Circ J. 2013;77:1307–14.

37. Ng KM, Lau YM, Dhandhania V, Cai ZJ, Lee YK, Lai WH, Tse HF and Siu CW. Empagliflozin Ammeliorates High Glucose Induced-Cardiac Dysfuntion in Human iPSC-Derived Cardiomyocytes. Sci Rep. 2018;8:14872.

38. Dambrot C, Braam SR, Tertoolen LG, Birket M, Atsma DE and Mummery CL. Serum supplemented culture medium masks hypertrophic phenotypes in human pluripotent stem cell derived cardiomyocytes. J Cell Mol Med. 2014;18:1509–18.

39. Woulfe KC, Ferrara C, Pioner JM, Mahaffey JH, Coppini R, Scellini B, Ferrantini C, Piroddi N, Tesi C, Poggesi C and Jeong M. A Novel Method of Isolating Myofibrils From Primary Cardiomyocyte Culture Suitable for Myofibril Mechanical Study. Front Cardiovasc Med. 2019;6:12.

40. Nemkov T, Reisz JA, Gehrke S, Hansen KC and D’Alessandro A. High-Throughput Metabolomics: Isocratic and Gradient Mass Spectrometry-Based Methods. Methods Mol Biol. 2019;1978:13–26.

41. Nemkov T, Hansen KC and D’Alessandro A. A three-minute method for high-throughput quantitative metabolomics and quantitative tracing experiments of central carbon and nitrogen pathways. Rapid Commun Mass Spectrom. 2017;31:663–673.

42. Sparagna GC, Johnson CA, McCune SA, Moore RL and Murphy RC. Quantitation of cardiolipin molecular species in spontaneously hypertensive heart failure rats using electrospray ionization mass spectrometry. J Lipid Res. 2005;46:1196–204.

43. Bligh EG and Dyer WJ. A rapid method of total lipid extraction and purification. Can J Biochem Physiol. 1959;37:911–917.

44. Yoon HR, Hong YM, Boriack RL and Bennett MJ. Effect of L-carnitine supplementation on cardiac carnitine palmitoyltransferase activities and plasma carnitine concentrations in adriamycin-treated rats. Pediatr Res. 2003;53:788–92.

45. FastQC [Internet]. 2019.

46. Dobin A, Davis CA, Schlesinger F, Drenkow J, Zaleski C, Jha S, Batut P, Chaisson M and Gingeras TR. STAR: ultrafast universal RNA-seq aligner. Bioinformatics. 2013;29:15–21.

47. Liao Y, Smyth GK and Shi W. featureCounts: an efficient general purpose program for assigning sequence reads to genomic features. Bioinformatics. 2014;30:923–30.

48. Liao Y, Smyth GK and Shi W. The R package Rsubread is easier, faster, cheaper and better for alignment and quantification of RNA sequencing reads. Nucleic Acids Res. 2019;47:e47.

49. Love MI, Huber W and Anders S. Moderated estimation of fold change and dispersion for RNA-seq data with DESeq2. Genome Biol. 2014;15:550.

50. Mi H, Muruganujan A, Ebert D, Huang X and Thomas PD. PANTHER version 14: more genomes, a new PANTHER GO-slim and improvements in enrichment analysis tools. Nucleic Acids Res. 2019;47:D419–D426.

51. Liao Y, Wang J, Jaehnig EJ, Shi Z and Zhang B. WebGestalt 2019: gene set analysis toolkit with revamped UIs and APIs. Nucleic Acids Res. 2019;47:W199–W205.

52. Oliveros JC. Venny. An interactive tool for comparing lists with Venn’s diagrams. 2007–2015.

53. Babicki S, Arndt D, Marcu A, Liang Y, Grant JR, Maciejewski A and Wishart DS. Heatmapper: web-enabled heat mapping for all. Nucleic Acids Res. 2016;44:W147–53.

