## Supplemental figures, methods, and tables for "Maturation of human induced pluripotent stem cell-derived cardiomyocytes for modeling hypertrophic cardiomyopathy"

### SUPPLEMENTAL MATERIAL

#### Supplemental Materials

Expanded Materials & Methods

Online Figures I-IX

Online Tables S1-S6

Online Video I-III Legends

References 40-53

### Expanded Materials and Methods

**hiPSC-CM Differentiation**: hiPSCs were induced into cardiomyocytes via modulation of Wnt signaling, according to previously established methods[5]. Specifically, hiPSC-CMs were plated into 24 well plates at a density of 375,000 cells per well. Cells were plated in mTESR1+Y-27632 (a ROCK inhibitor, Cayman Chemicals, Ann Arbor, MI), and were then cultured in mTESR1 (STEMCELL Technologies, Vancouver, BC, CA) for four days with daily media changes. After four days, media was changed to RPMI (RPMI) with B-27™ supplement without insulin (B-27–ins; 1:50, Life Technologies, Carlsbad, CA), and 8μm CHIR99021 (Cayman Chemicals, Ann Arbor, MI), a GSK-3β inhibitor. This was considered day 0 (D0) in terms of cardiomyocyte age. Exactly 24 hours later, media was replaced with RPMI/B-27-ins. 2 days later (D3), media was changed again: at this point, old media (from D1) was combined with fresh RPMI/B-27-ins, in a 1:1 ratio, supplemented with 5μM IWP2 (Thermo Fisher Scientific, Waltham, MA), a Wnt inhibitor. 2 days later (D5), media was changed to RPMI/B-27-ins. On D7, media was replaced with RPMI with B-27 supplement with insulin (B27, 1:50, Life Technologies, Carlsbad, CA). Cells were then maintained with RPMI/B27 with media changes every 3 days. Contracting cells were typically observed on D10-12 post induction, while large areas of plates typically began beating by D15-18.

Once robust cellular contraction was observed in most plates, cells were digested using 0.25% Trypsin-EDTA (Thermo Fisher Scientific, Waltham, MA), pooled, and replated into 6 well plates at a ratio of 1-24 well plate well to 1-6 well plate well, in RPMI medium supplemented with Hyclone™ 20% FBS (GE Healthcare, Chicago, IL) and Y-

27632. Media was changed after 48 hours, to RPMI/B27. On D20-25 post induction, cardiomyocytes were selected for using the lactate method[6]: cells were washed with PBS, then media was changed to glucose-free DMEM supplemented with 4mM lactate. Cells were cultured in lactate media for 5 days, with media changed on the third day of lactate culture. Media was then changed again to RPMI/B27. In the case of cells cultured under MM or MPAT conditions, media was then changed to MM after 3 days. Switching medium directly from lactate to MM appeared to cause significant levels of cell death so was avoided.

**Patterned surface preparation:** Patterned surfaces were created by drawing 20 micro lapping paper across 22mmx22mm plastic coverslips (VWR International, Radnor, PA, Cat. No. 48376-049), 20-30 times per coverslip. Coverslips were held in place during patterning to ensure consistency of direction of the grooves created by the patterning process. Surfaces were then washed in soapy water to remove plastic dust or other contaminants, and soaked repeatedly in ultrapure water. For imaging studies, coverslips were cut into 11*11mm squares. To sterilize, surfaces were incubated overnight in 100% ethanol. The following day, surfaces were placed in 6 well plates, and residual ethanol allowed to evaporate. Surfaces were then incubated under cell culture ultraviolet for 10-15 minutes for sterilization. Fibronectin solution in PBS (6µg/mL, Corning Inc, Corning, NY) was then added to each surface, and fibronectin coating allowed to occur at room temperature for 1-2 hours. Surfaces were then washed in PBS, and allowed to completely dry before adding cells to surfaces. For imaging studies, GLUC and MM hiPSC-CMs were cultured on unpatterned surfaces which were prepared in an identical fashion to patterned surfaces except for the lapping paper step. For other experiments, GLUC and MM cells were cultured on fribronectin-coated 6 well plate wells.

**Cardiomyocyte replating onto surfaces:** On day 35-40 post induction, cardiomyocytes were replated onto patterned surfaces. Cells were digested for 7-9 minutes using Accumax (Innovative Cell Technologies, San Diego, CA) at 37°C, then digestion was halted by addition of RPMI-20. Cells were counted, then pelleted by centrifuging at 1000g for 3 minutes, then resuspended in RPMI-20 supplemented with Y-27632. When plating onto large (2cmx2cm) patterned surfaces, 600-750,000 cardiomyocytes were plated per surface in 500-750µL cultured medium, which was sufficient to create a confluent, uniformly beating cell sheet. When plating onto small (8mmx8mm) patterned or unpatterned surfaces for imaging studies, 15,000 cells were plated per surface. Cell suspension was added dropwise onto each surface; cell medium stayed on the surfaces via surface tension. (This step needs to be conducted very carefully, as it is easy to cause cell solution to run off of surface: it is also essential that the surface be completely dry before adding cells). Cells were then allowed to attach to surface at 37°C for 45 minutes before adding an additional 1.5mL medium. Medium was changed to MM after 48 hours, and then changed biweekly thereafter.

**Cell imaging studies**: Cells on small unpatterned or patterned surfaces in 24 well plates were used for immunofluorescent staining. Briefly, cells were washed with PBS, then fixed either with 4% paraformaldehyde (PFA) in PBS (10 minutes at room temperature) or a 1:1 mixture of methanol and acetone (5minutes at -20°C). After fixation, cells were washed with PBS, then permeabilized with 0.2% Triton-X 100 in PBS (10 minutes at room temperature). Methanol/acetone fixation was used for images shown in figure 1 (staining for α-actinin, MYBPC3, c-TNI, and MLC-2V). PFA fixation was used for all other experiments. All measurements of cell area and hypertrophy were conducted on PFA-fixed cells.

For immunofluorescent staining, after permeabilization, cells were treated with blocking buffer (10% horse serum in PBS) for 30 minutes at room temperature. Cells were then treated with primary antibodies diluted in 5% horse serum/PBS for 2 hours, washed four times in PBS for 5 minutes, and treated with secondary antibodies in 5% horse serum/PBS for 1 hour, washed twice with PBS for 5 minutes, counterstained with Hoescht 3342 in PBS for 5 minutes, and washed twice with PBS for 5 minutes, all at room temperature.

For cell area measurements, cells were PFA fixed and stained for α-actinin as described above, then counterstained with HCS CellMask™ Orange (Thermo Fisher Scientific, Waltham, MA), diluted 1:2,500 in PBS, for 30 minutes at room temperature.

For B-type natriuretic peptide (BNP) staining, an adaption of a previously published method was used[27]: cells were PFA fixed, then blocked and permeabilized for 30 minutes with 5% milk/0.1% Triton X-100/PBS (blocking solution) for 30 minutes at room temperature. Cells were then incubated with pro-BNP antibody diluted in blocking solution overnight at 4°C. Cells were then washed four times in 0.05% Triton-X 100/PBS for 5 minutes, and incubated with secondary antibody diluted in blocking solution for 1 hour at room temperature. Cells were then washed twice in 0.05% Triton-X 100/PBS for

5 minutes, counterstained with Hoescht 3342 as described above, and washed twice more.

All imaging was conducted on an EVOS FL Cell Imaging System (Thermo Fisher Scientific, Waltham, MA). All image analysis including measurements of cell area, circularity, and sarcomere length were conducted using Image J.

For immunostaining, the following primary antibodies were used: **1)** Ventricular myosin light chain 2 (MLC-2v): Proteintech 10906-1AP rabbit antibody at 1:400 (Proteintech Group, Rosemont, IL) ; **2)** Alpha actinin: Sigma-Aldrich A7811 mouse antibody at 1:400 (MilliporeSigma, Burlington, MA) ; **3)** Myosin binding protein C3: Santa Cruz E-7 sc-137180 mouse antibody at 1:400 (Santa Cruz Biotechnology, Inc., Dallas, TX); **4)** Cardiac troponin I: Phosphosolutions 2010-TNI rabbit antibody at 1:200 (PhosphoSolutions, Aurora, CO); **5)** B-type natriuretic peptide (and pro-nt-BNP): Abcam ab13115 [15F11] mouse antibody at 1:200 (Abcam, Cambridge, UK). The following secondary antibodies were used: **1)** Alexa Fluor 488 goat anti-rabbit A11034; **2)** Alexa Fluor 546 goat anti-rabbit A21428; **3)** Alexa Fluor 488 A11029 goat anti-mouse; **4)** Alexa Fluor goat anti-mouse 546 A21422 (Thermo Fisher Scientific, Waltham, MA). All secondary antibodies were used at 1:400 dilution.

**MYH6/7 measurements**: Prior to lysis, cells were detached by treatment with Accumax, counted, and pelleted. Pellets were then lysed by resuspending pellets in isoelectric focusing buffer (IEF) with protease inhibitors. For human and mouse heart samples, approximately 20mg of powdered left ventricular tissue per sample was resuspended in IEF by vortexing, then spun at 5000g for 5 minutes at 4°C to remove insoluble tissue. To determine protein content, a 10μL aliquot of each lysate (diluted as necessary) was pH neutralized by the addition of 40μL 0.12N HCl, then protein concentration was measured by Bradford Assay on these neutralized samples and compared to a standard curve to calculate protein concentrations. 10μg of protein per sample was loaded onto a degassed 6% SDS-PAGE gel with the following modifications: resolving gel buffer pH at 9.0; running gel buffer pH at 8.2; inner gel buffer supplemented with β-mercaptoethanol at 600 μL/L; and separating acrylamide/bis ratio 1:100. These gels were run overnight at 0.12V at 4°C, and stained with modified Coomassie blue/silver solution to stain protein.

**Immunoblotting:** Prior to lysis, cells were detached by treatment with Accumax, counted, and pelleted. Pellets were then lysed by resuspending pellets in ice-cold lysis buffer (150 mM NaCl, 50 mM Tris-Cl pH 7.4, 1 mM EDTA, 1% Triton, with Complete mini tablet (Roche, Basel, CH), and 1 mM phenylmethylsulphonyl fluoride freshly added before use). Protein concentrations were determined using a BCA assay according to manufacturer's instructions (Bio-Rad Laboratories, Hercules, CA). 4μg of protein per sample was loaded onto a precast 4-12% Criterion Tris-HCL gel (Bio-Rad Laboratories, Hercules, CA). Protein was transferred overnight onto activated polyvinylidene fluoride (PVDF) membrane overnight at 0.12V at 4°C. For Western blotting, membranes were blocked for 1 hour with 5% milk/TBST buffer for 1 hour at room temperature, incubated with primary antibody in TBST buffer overnight at 4°C, and incubated in secondary antibody in 5% milk/TBST buffer for 1 hour at room temperature, with TBST washes between antibodies. The following primary antibodies were used for Western blotting: **1)** Ventricular myosin light chain 2 (MLC-2v): Proteintech 10906-1AP rabbit antibody at

1:1000; **2)** Atrial myosin light chain 2 (MLC-2a): Proteintech 17283-1-AP rabbit antibody at 1:1000 (Proteintech Group, Rosemont, IL); **3)** Myosin binding protein C3: Epitomics 3153-1 rabbit antibody at 1:1500 (Epitomics, Inc, Burlingame, CA); **4)** Alpha actinin: Sigma A7811 mouse antibody at 1:2000 (MilliporeSigma, Burlington, MA); **5)** Cardiac/slow skeletal troponin I: Abcam Ab10239 [MF4] at 1:1:1000 (Abcam, Cambridge, UK, note that while this antibody is listed as cardiac specific, the epitope it was raised against is common to cardiac and slow skeletal TNI); **6)** Non-muscle myosin (Myosin IIB): Sigma M7939 rabbit antibody at 1:1000 (MilliporeSigma, Burlington, MA); **7)** alpha tubulin: Cell Signaling #3873 mouse antibody at 1:5000 (Cell Signaling Technology, Danvers, MA). The following secondary antibodies were used: **1)** For mouse primary antibodies: Southern Biotech goat anti-mouse 1031-05, at 1:2000-1:5000 (for alpha tubulin, was used at 1:5000, Southern Biotech, Birmingham, AL); **2)** For rabbit primary antibodies: Sigma A9169 at 1:2000-1:5000 (MilliporeSigma, Burlington, MA). Membranes were developed using Supersignal™ Pico Plus (Thermo Fisher Scientific, Waltham, MA).

**Metabolomic Screening:** Prior to sample submission, cells were detached by treatment with Accumax, counted, and pelleted, and flash frozen using liquid nitrogen. 131,000-875,000 cells were submitted per individual sample, with samples submitted from four independent hiPSC-CM inductions. Metabolites from frozen cell pellets were extracted using ice-cold methanol/acetonitrile/water (5:3:2) at a ratio of 2e6 cells per mL by vortexing for 30 minutes at 4°C. Samples were clarified through centrifugation (10 minutes at 10,000 rpm, 4°C) and 10 µL of supernatant was analyzed using a 5 minute C18 gradient on a Thermo Vanquish UHPLC coupled online to a Thermo Q Exactive mass spectrometer operating in positive and negative ion modes (separate runs) as previously described in detail[40]. Metabolites were assigned using Maven (Princeton University) in conjunction with the KEGG database. Assignments and quality control were performed as previously described[41].

**Cardiolipin Quantification:** Cardiolipin was quantified using previously published methods with normal phase liquid chromatography coupled to electrospray ionization mass spectrometry in an API 4000 mass spectrometer (Sciex, Framingham, MA) [42]. To perform this assay with hiPSC-CMs, cells were dissociated using Accumax and counted, with1-7.5 x$10^5$ cells from each condition were harvested for the assay. Cells were flash frozen in liquid nitrogen prior to use. Lipids were extracted using a modified Bligh Dyer method according to previously published methods with 1000 nmoles tetramyristal cardiolipin as an internal standard (Avanti Polar Lipids, Alabaster, AL) [42, 43].

**Carnitine Palmitoyl Transferase Activity Assay:** Rates of CPT1 and CPT2 were quantified using a 14C carnitine based radioactive assay[44]. The assay measures the activity of CPT1 by permeabilizing the plasma membrane and measuring the production of palmitoyl carnitine from palmitoyl CoA. By permeabilizing the mitochondrial inner membrane and adding malonyl CoA to inhibit CPT1, the activity of CPT2 was measured. To perform this assay with iPSC cells, cells were dissociated using Accumax and counted, with 1.75-7.5 x$10^5$ cells from each condition were harvested for the assay. CPT activity was normalized to cell counts.

**RNA extraction and cDNA generation:** RNA was generated via a column-free RNA extraction method. Cells were initially detached and counted with Accumax, and were then lysed in 1mL TRIzol™ (Thermo Fisher Scientific, Waltham, MA) reagent per cell pellet. 250μL chloroform was then added to each sample, mixed by vortexing, and centrifuged at 12,500rpm for 15 minutes at 4°C. Upper phase was then decanted, and an equal volume of isopropanol and 2μL glycogen to each RNA solution. Samples were centrifuged again at 12,500rpm for 20 minutes at 4°C, and supernatants decanted from RNA pellet. Pellets were washed with 70% ethanol/ultrapure DNAase/RNAase-free $H_2O$, and centrifuged again at 12,500rpm for 5 minutes at 4°C. RNA pellets were then resuspended in 17μL ultrapure DNAase/RNAase-free $H_2O$ per pellet.

Genomic DNA was removed via DNAase I treatment. RNA was then recrystallized via addition of a 100mM $LiCl_2$/74% (vol:vol) EtOH/2% (vol:vol) glycogen solution to the RNA solution. This solution was then incubated at -80°C for 90 minutes; RNA was then pelleted by centrifugation at 12,500rpm for 30 minutes at 4°C. Pellets were then washed once with an ice cold 70% (vol:vol) ethanol solution, and centrifuged again at 12,500rpm, and resuspended in 20μL RNAase/DNAase-free $H_2O$. RNA yield was determined using the Nanodrop™ (Thermo Fisher Scientific, Waltham, MA). cDNA was generated using the iScript cDNA Kit (Bio-Rad) according to manufacturer's instruction, using 250ng-1μg RNA as substrate, using a mixture of random primers. qPCR gene expression analysis was performed using SYBR Green Supermix (Bio-Rad), according to manufacturer's instructions, using 0.5μL cDNA in a 20μL total reaction volume. Each reaction was performed in duplicate.

The following primer sets were used for qPCR analysis:

| | | Primer sequence |
|---|---|---|
| Human ***TNNI3*** | for | 5'- TGC TTC ACA GTG GAG CTG ATA -3' |
| | rev | 5'- GCT GCA ATA TGC AAT GGA GTG -3' |
| Human ***MYL2*** | for | 5'- CGC CAA CTC CAA CGT GTT CT -3' |
| | rev | 5'- CCA TCC CTG TTC TGG TCC AT -3' |
| Human ***ACTN2*** | for | 5'- CTG CTG CTT TGG TGT CAG AG -3' |
| | rev | 5'- TTC CTA TGG GGT CAT CCT TG -3' |
| Human ***NPPA*** | for | 5'- GAC TCC TCT GAT CGA TCT GC -3' |
| | rev | 5'- GTT ATC TTC AGT ACC GGA AGC -3' |
| Human ***NPPB*** | for | 5'- TCC TGC TCT TCT TGC ATC TGG -3' |
| | rev | 5'- TTT GCC CTG CAA ATG GTT G -3' |
| Human ***ACADVL*** | for | 5'- CCT GGA AGG TGA CAG ATG AAT G-3' |
| | rev | 5'- CCC TCA AAG ATC CGG AAG ATG -3' |
| Human ***CPT1A*** | for | 5'- CAC ACA CAG TGG AAT GGA AAT G -3' |
| | rev | 5'- CCT GCT TGG ATG ATG CTA AAT G -3' |
| Human ***CPT1B*** | for | 5'- GCT ATG TGT ATC CGC CTT CTA TC -3' |
| | rev | 5'- GAC TCT AGG TAC CGC TGA ATT G -3' |
| Human ***CPT2*** | for | 5'- GCT TTG ACC GAC ACT TGT TTG -3' |
| | rev | 5'- TGT GGT TTA TCT GCC CGT ATG -3' |
| Human ***TAZ*** | for | 5'-GAG AAC AAG TCG GCT GTG G-3' |
| | rev | 5'-GCT GGA GGT GGT TGT GGA -3' |
| Human ***18S rRNA*** | for | 5'- GCC GCT AGA GGT GAA ATT CTT A -3' |
| | rev | 5'- CTT TCG CTC TGG TCC GTC TT -3' |

**RNA-seq protocol:** Total RNA was prepared as described above. Prior to submission, RNA quality was confirmed via analysis on Tapestation (Agilent Technologies, Santa Clara, CA). Library preparation and RNA-seq was performed by BGI Genomics (Shenzhen, CN). Briefly, mRNA was purified from total RNA using oligo(dT) magnetic beads, then fragmented. First strand cDNA was generated using random hexamers, followed by second strand synthesis. cDNA was then subjected end repair, and 3'-adenylation, adapter ligation, PCR amplification, and circularization, creating a single strand circle DNA library. RNA-seq was performed at a read depth of 40 million reads, using paired-end sequencing, on the DNBSeq Platform.

Quality control of the RNA-seq data was performed with FastQC[45]. The RNA-seq reads were mapped to human GRCh38 reference genome with the splice-aware STAR aligner v2.7[46]. The read counts were generated from the aligned reads with featureCounts[47] function in the RsubRead package[48]. The read counts were normalized with DESeq2 R package[49].

**Gene Ontology and Pathway Analysis:** Gene ontology analysis was performed using PANTHER and KEGG. Prior to analysis, RNA-seq data was filtered to only include to top 15,000 most highly expressed genes by total expression (across the GLUC, MM and MPAT conditions). Data were then sorted by fold change between each 2 condition comparison. These comparisons were filtered for genes showing either a 1.5X upregulation or downregulation between groups in question, and these filtered data fed separately into PANTHER for gene ontology analysis[50], or WebGestalt for KEGG Pathway analysis[51]. For Panther, 'Statistical overrepresentation test' was selected, using 'GO biological process, complete', and the Homo sapiens whole-genome list as reference. For WebGestalt, 'Overrepresentation analysis' was also selected, using 'pathway/KEGG', a minimum of 10 genes per category, and the Homo sapiens whole genome list as reference.

**Cardiomyocyte Isolation**: 15 minutes prior to beginning isolation, mice were injected with 300μL of heparin (10,000 units/mL). Mice were then anesthetized via inhalation of isofluorane, and when becoming unresponsive to toe pinch were killed via cervical dislocation. The chest was then opened, and the heart was removed rapidly and placed in ice-cold perfusion buffer (10 mM creatine monohydrate, 30 mM Taurine, 5.6 mM D-glucose, 4.6 mM $NaHCO_3$, 10 mM BDM, 120 mM NaCl, 15 mM KCl, 0.6 mM $Na2HPO_4$, 0.6 mM $KH2PO_4$, 1.2 mM $MgSO_4$, 10 mM Hepes, filtered at 0.40 μm, pH 7.4). The heart was washed briefly in perfusion buffer, and excess fat/thymus tissue trimmed off. The aorta was then cannulated using a blunted 20-gauge needle, and was mounted on a Langendorff apparatus (Radnoti, Covina, CA) with a heating pump kept at 37°C for the duration of the isolation (VWR International, Radnor, PA). Hearts were mounted on the Langendorff apparatus within 3-7 minutes of removing from the body. The heart was then perfused in retrograde fashion at a rate of 4mL/minute, driven by a Rainin™ Rabbit peristaltic pump (Mettler Toledo, Columbus, OH). To remove blood from the heart's vessels, it was first perfused for 3 min using perfusion buffer. Buffer was then switched to calcium-free digestion buffer (perfusion buffer with 1.3 mg/mL collagenase II) for 3 min and was then switched to digestion buffer supplemented with 28 nM $CaCl^2$. Hearts were perfused with high-calcium digestion buffer for another 6 to 10 minutes, once the heart became extremely soft and compliant. The heart was then removed from the cannulating needle and placed in a 60mm culture dish with 2.5 mL high-calcium digestion buffer. The atria and right ventricle were then removed, and the left ventricle dissociated using a pipette and forceps. Once the heart was completely dissociated, digestion was halted by the addition of 7.5 mL stopping buffer (perfusion buffer with 10% (vol/vol) FBS and 12.5

nM $CaCl_2$). The cardiomyocyte suspension was then filtered through 200-μm mesh into a 50-mL conical tube, and the culture dish washed with an additional 10mL stopping buffer, which was then filtered into the conical tube as wlel. This cell suspension was then allowed to incubate for 10 minutes at room temperature, allowing a cardiomyocyte-rich pellet to form. The fibroblast-rich supernatant was discarded, while the pellet with was resuspended in 10 mL of stopping buffer. 100mM $CaCl_2$ solution was then added to this buffer in a stepwise manner, with 2-min intervals between each addition: calcium concentration was first increased to 112.5 μmol/L, then to 312.5 μmol/L, then 712.5 μmol/L, and finally to 1.4 mmol/L. The cell suspension was then centrifuged at 1,000 × g for 3 min at room temperature. Cells were then inspected and counted, and if <60% of myocytes were rod-shaped (viable) the cells were not used for experimentation. Cardiomyocytes were then resuspended in plating medium [perfusion buffer with 2.5% (vol/vol) FBS, 1% (vol/vol) penicillin/streptomycin (P/S), and 1.4 mM $CaCl_2$, and plated on either 6 well plates (for myofibril mechanics) or 24 well plates (for hypertrophy imaging) coated with 9–10 μg/mL Laminin. Cells were incubated in plating buffer for 1-2 hours at room temperature to promote attachment; myocytes then were washed 2X in sterile PBS to remove dead cells, and media was changed to myocyte culture medium [MEM with 0.2% (wt/vol) BSA, 10 mM Hepes, 4 mM $NaHCO_3$ 10 mM creatine, 0.5% (vol/vol) insulin-selenium-transferrin (Thermo Fisher Scientific, Waltham, MA), 1% (vol/vol) penicillin/streptomycin, 1μM Blebbistatin (Cayman Chemicals, Ann Arbor, MI), filtered at 0.22 μm pH brought to 7.4, for 30-60 minutes before treatment.

**Myofibril mechanics:** Myofibril mechanics were performed on myofibrils isolated from human adult cardiac tissue, mouse ventricular cardiomyocytes, and stem cell derived cardiomyocytes. To isolate myofibrils from adult cardiac tissue, frozen heart sections were cut into ~5mm strips and skinned overnight in 0.5% Triton-X in rigor solution (132 mM NaCl, 5 mM KCl, 1 mM MgCl2, 10 mM Tris,5 mM EGTA, pH 7.1) containing protease inhibitors (10 µM leupeptin, 5 µM pepstatin, 200 µM PMSF and 10 µM E64), as well as 500 µM $NaN^3$ and 500 µM DTT at 4°C. These skinned LV strips were then washed several times in relaxing solution and homogenized (Tissue-Tearor, Thomas Scientific, Swedesboro, NJ) in relaxing solution (pCa 9.0) containing protease inhibitors.

To isolate myofibrils from adult mouse cardiomyocytes and stem cell derived cardiomyocytes, cells were lysed and myofibrils prepared using an adaption from a recently described method[39]. In brief, cells were lysed in relaxing solution with protease inhibitors and 20% (w/vol) sucrose, using a scraper. The lysed cell solution was then transferred into a microfuge tube and incubated on ice for 10 minutes with periodic vortexing. The cell/myofibril solution was then centrifuged at 1,500g for 5 minutes at 4°C, and the cell/myofibril pellet resuspended in fresh relaxing solution with protease inhibitors. This process was repeated twice to remove any residual sucrose, and the cell/myofibril solution was then homogenized. A significantly more strenuous homogenization protocol was required for AMVCMs compared to hiPSC-CMs.

Myofibril mechanics were quantified using the fast solution switching technique on a custom-built experimental rig. Myofibril suspensions were plated onto a temperature controlled glass coverslip (15°C) containing relaxing solution. Single myofibrils and myofibril bundles were then picked up from the surface of this coverslip and mounted between two glass micro-tools. One tool, the stretcher, was linked to a motor which could produce rapid changes in length (Mad City Labs, Madison, WI), while the second tool was a calibrated cantilevered force probe (6-8 μm/μN; frequency response 2-5 KHz), which was colored with black ink to increase probe sensitivity. After mounting myofibrils between microtools, they were stretched to 5-10% above slack myofibril length. Average sarcomere lengths and myofibril diameters were measured using ImageJ software. Mounted myofibrils were then activated and relaxed by rapidly changing the position of a double-barreled pipette simultaneously releasing activating solution (pCa 4.5) and relaxing solution (pCa 9.0) from each barrel. By rapidly changing the position of the pipette, the mounted myofibril could be induced to activate and relax. From the activation/relaxation trace produced, the following parameters were acquired: resting tension, the passive tension produced in relaxing conditions, induced by a push on the stretcher; maximum tension (mN/mm2), the greatest level of active tension generated by the myofibril at full calcium activation (pCa 4.5); activation kinetics ($k_{Act}$), the rate constant of tension development following myofibril activation; reactivation kinetics ($k_{Tr}$), the rate constant of tension redevelopment following myofibril release/stretch to break cross bridges; slow phase relaxation kinetics ($k_{REL,SLOW}$), the kinetics of the initial slow/linear phase of myofibril relaxation, slow phase relaxation time ($T_{LIN}$), the duration of the slow phase of relaxation, from the switching of the pipette to relaxing solution to the beginning of the fast phase of relaxation; fast phase relaxation kinetics ($k_{REL,FAST}$), the rate constant of the fast (exponential) phase of relaxation.

**Statistical Analysis:** All data was first tested for normality. Data which did not initially demonstrate normality was then log2 transformed. Metabolomics data was log10 transformed instead, because log2 transformation did not achieve normality. For data which demonstrated normality (either before or after transformation), for two-group comparisons, an unpaired t-test was used. For multi-group comparisons, a one-way ANOVA with Tukey's multiple comparison test was used. For data which did not demonstrate normality after log transformation, for two group comparisons, a Mann-Whitney test was used. For multi-group comparisons, a Kruskal-Wallis test followed by Dunn's multiple comparison test was used.

P values or adjusted P values of less than 0.05 were considered to be significant. All data is displayed as mean plus or minus standard error the mean (SEM).

Venn's Diagrams were created using Venny 2.1[52]. Heatmaps were created Heatmapper[53]; samples were clustered by centroid linkage. All other statistical analysis was performed using GraphPad Prism 8.3.0 (GraphPad Software, San Diego, CA).

Supplemental Figure I

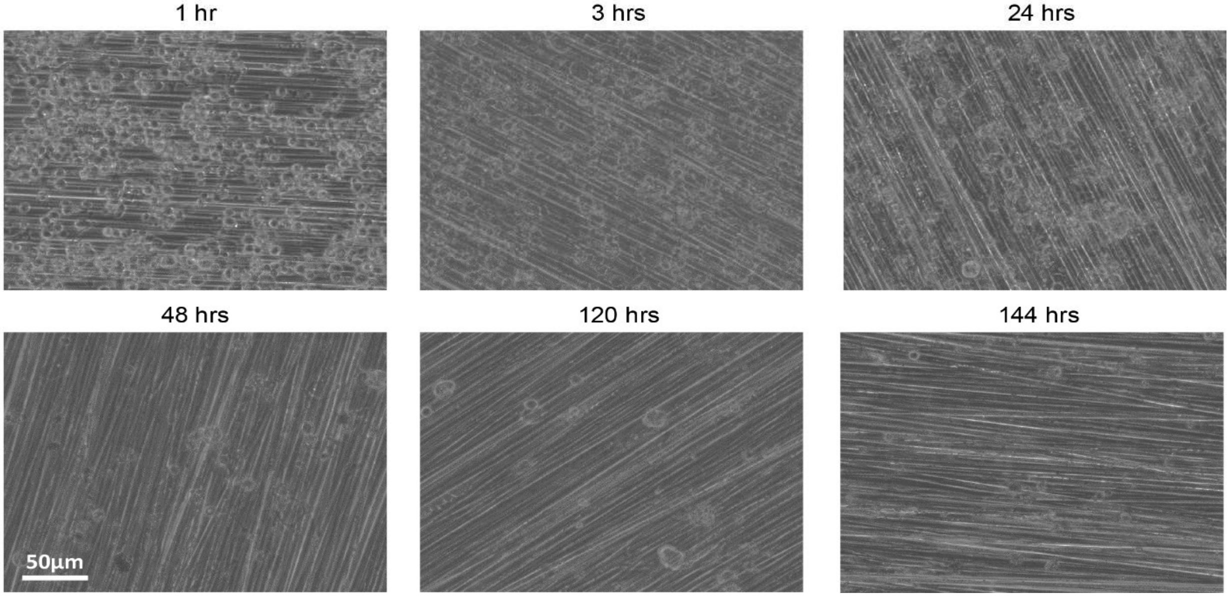


**Supplementary Figure Legends**

**Figure S1. Attachment of hiPSC-CMs to patterned surfaces.** Brightfield images of cells at various time points after plating. Cells can be seen to begin to attach and align to grooves within 3 hours of plating, with the majority of cells having aligned within 24 hours, and nearly all cells after 48 hours. After several days of plating, cells plated at high density form a sheet with coordinated contraction along the direction of micropatterning (also see **Video 3**).

Supplemental Figure II

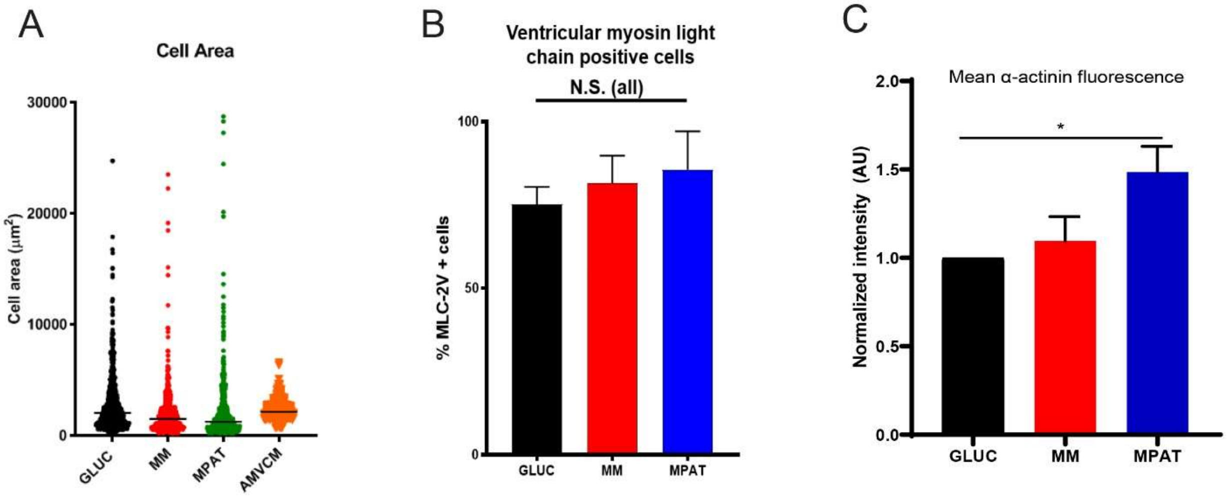


**Figure S2. Fatty acid medium and patterning improves hiPSC-CM maturity. A,** Comparison of cell area distribution of hiPSC-CMs cultured under each condition, and adult mouse ventricular cardiomyocytes. Data pooled from 507-706 cardiomyocytes per condition from 4-5 independent experiments. **B,** Percentage of cells positive for ventricular myosin light chain (MLC-2V) in each culture condition. Data from 3 independent experiments, 75-325 cells assessed per condition per experiment. **C,** Mean α-actinin staining fluorescence intensity per cell, normalized to glucose-cultured hiPSC-CMs. Data from 5 independent experiments, 100-150 cells assessed per condition per experiment. *P< 0.05, one way ANOVA with Tukey's multiple comparison test. N.S., nonsignificant. Data are mean + SEM.

Supplemental Figure III

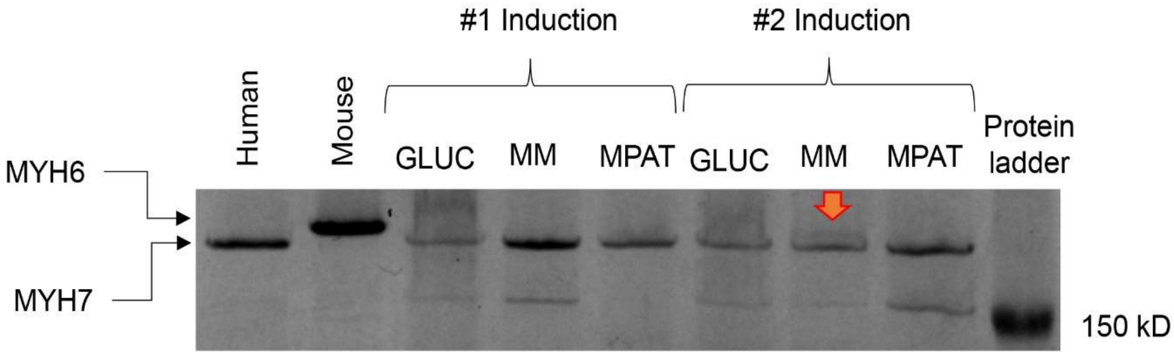


**Figure S3. hiPSC-CM protein expression and myofibril mechanics.** Blue silver staining of myosin heavy chains (MYH6 and 7, indicated by arrows) in hiPSC-CMs, as well as from a donor human heart lysate (as a positive control for MYH7) and a 12 week old mouse heart (positive control for MYH6). In a single MM hiPSC-CM sample, significant MYH6 could be detected (indicated by thick arrow); all other samples appear to express exclusively MYH7.

Supplemental Figure IV

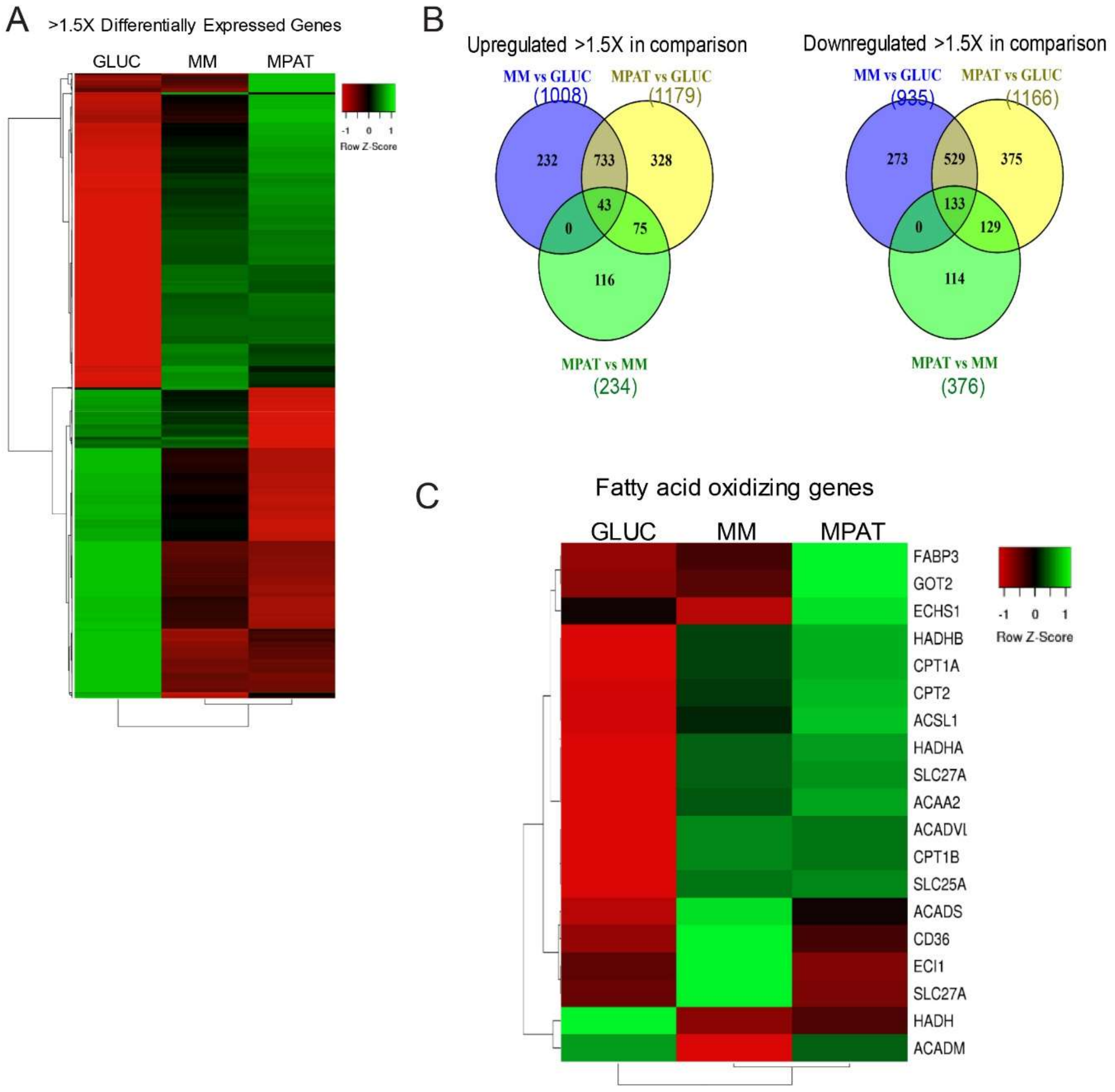


**Figure S4. RNA-seq on hiPSC-CMs cultured under each condition. A,** Heatmap of all genes with >1.5 fold change (increase or decrease in expression) between GLUC and MPAT cells. **B,** Diagram of overlap in genes displaying >1.5X increase (left panel) or decrease (right panel) in expression in indicated comparison. Many of the same genes are upregulated or downregulated comparing GLUC versus MM and GLUC versus MPAT, whereas other comparisons have relatively little overlap. **C,** Heatmap of selected genes involved in fatty acid oxidation. Unsupervised hierarchical clustering performed on both rows and columns, by centroid linkage.

Supplemental Figure V

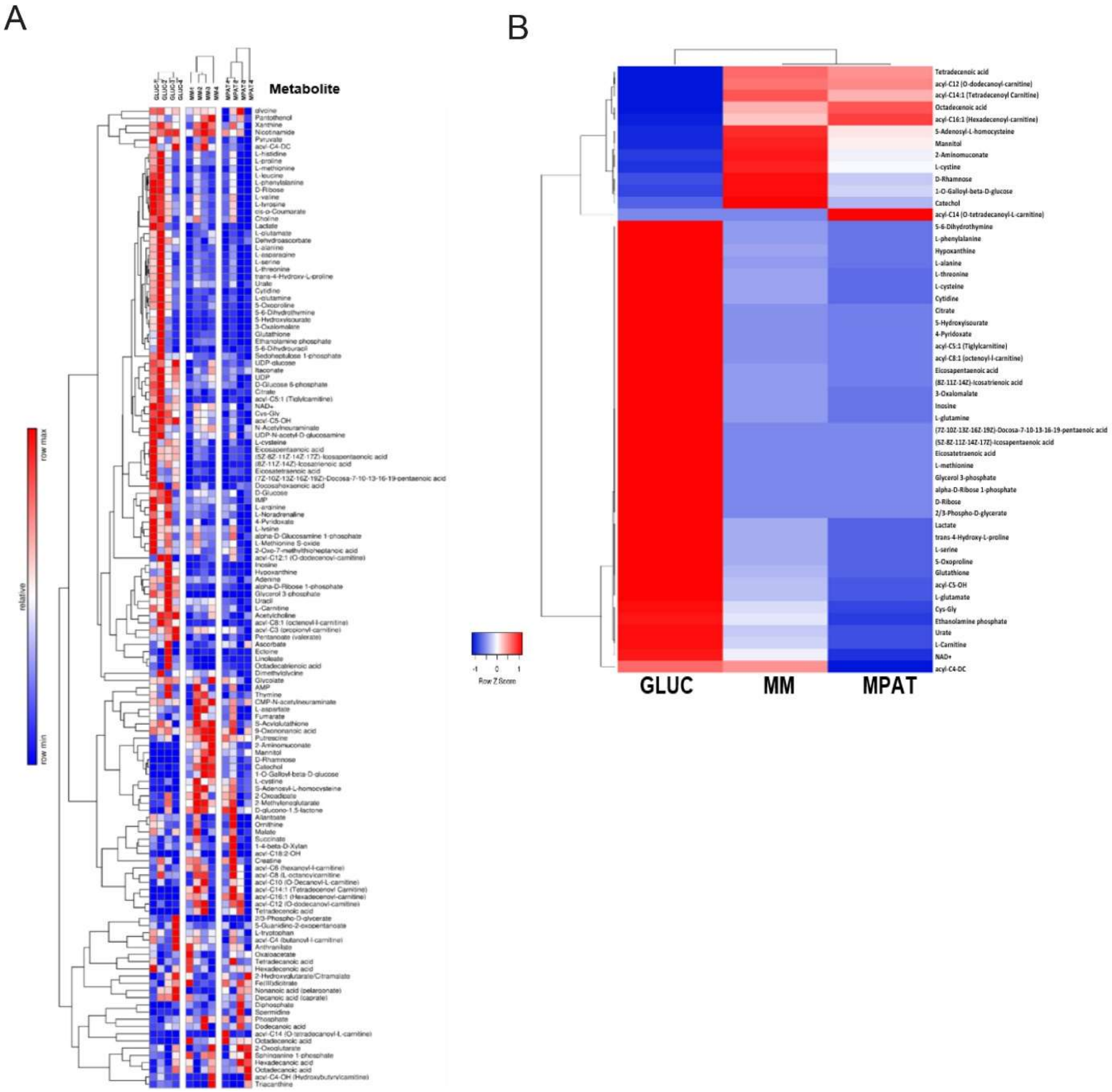


**Figure S5. Maturation methods induce broad changes in hiPSC-CM metabolites. A,** Heatmap of all polar metabolites detected in screening, showing individual experimental replicates from each separate cardiomyocyte induction. **B,** Heatmap of significantly differentially expressed metabolites, as assessed by one way ANOVA and Tukey's post test on log10 transformed data. Unsupervised hierarchical clustering performed on both rows and columns, by centroid linkage.

Supplemental Figure VI

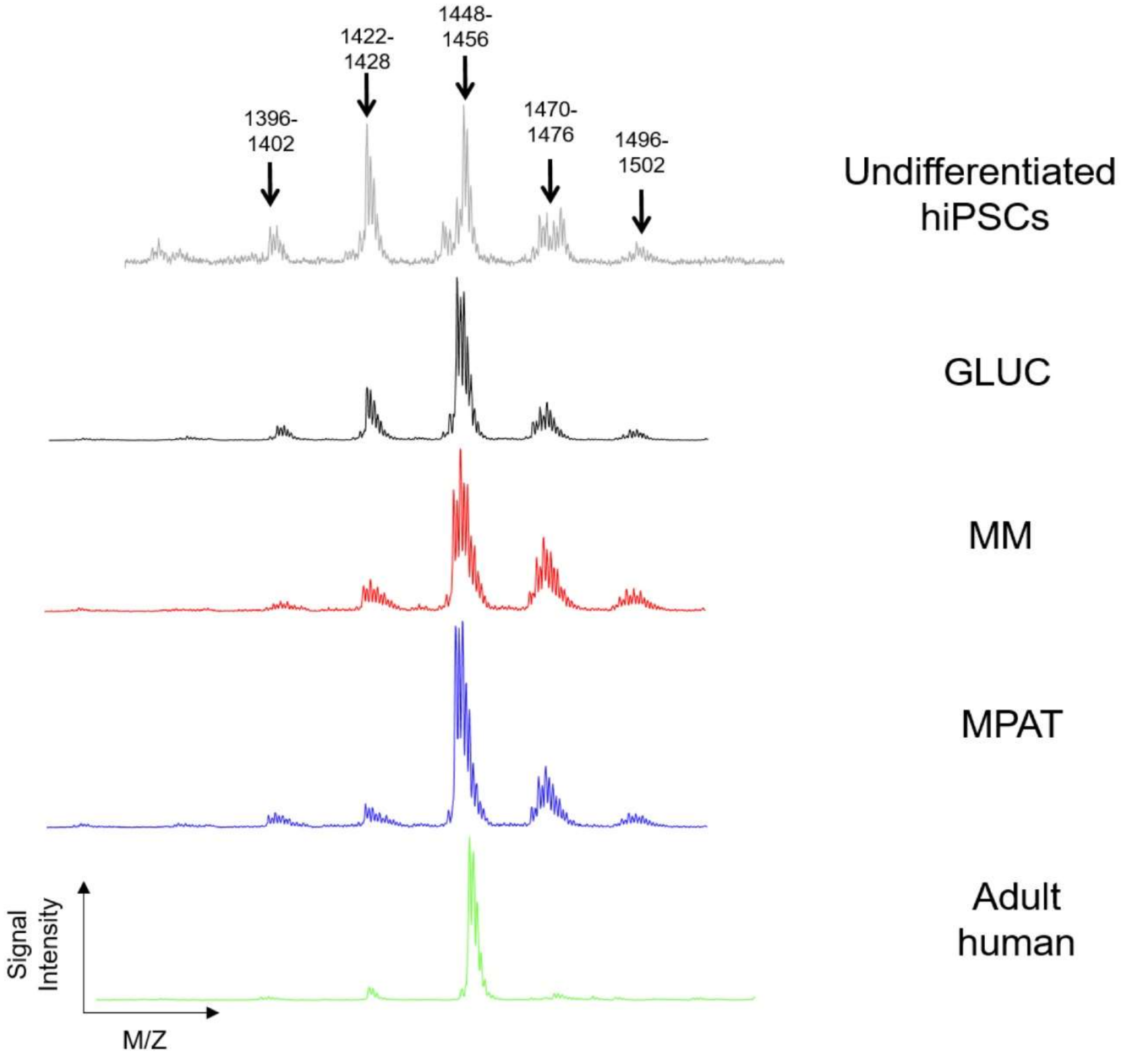


**Figure S6. Maturation methods induce changes in cardiolipin side chain content in hiPSC-CMs.** Representative CLs spectra from hiPSCs and hiPSC-CMs cultured under each condition, as well as from a donor heart. Peaks are labeled by total MW range of CLs contained within peak.

Supplemental Figure VII

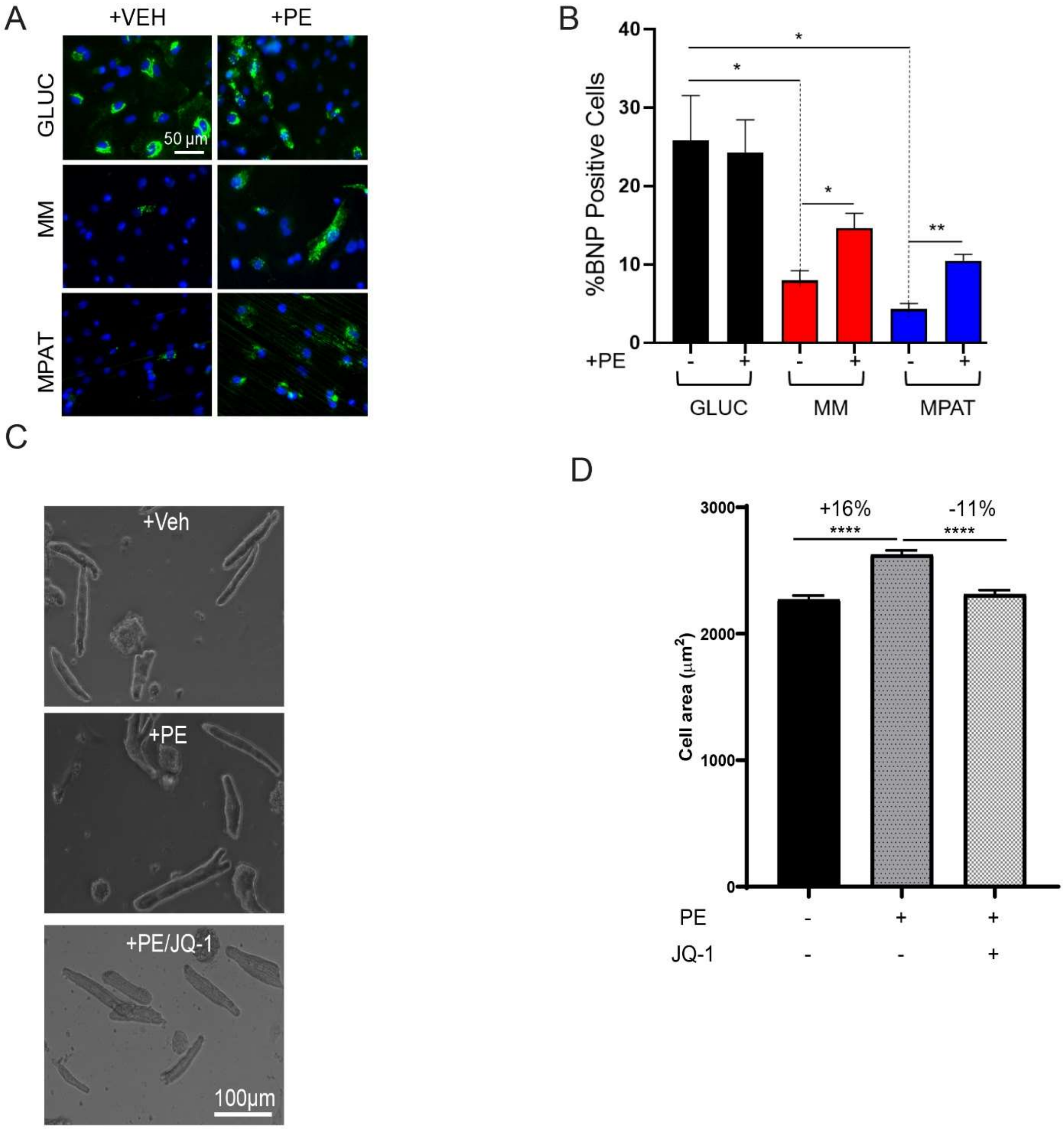


**Figure S7. Hypertrophic response in hiPSC-CMs and adult mouse ventricular cardiomyocytes (AMVCMs)**. **A-B,** Representative images of BNP staining and quantification of BNP and pro-NT-BNP-positive hiPSC-CMs cultured and treated as described in **Figure 5A-B**. 500 or more cells were analyzed per experiment in cells from

3 independent inductions. **C-D,** Representative brightfield images and cell area of AMVCMs under indicated conditions. AMVCMs were isolated from 8-12 week old adult mice then treated with vehicle, 10 µmol/L phenylephrine (PE) or 10 µmol/L PE and 1 µmol/L JQ1 for 72 hours, then fixed. Cell areas measured on 664-706 cardiomyocytes pooled from four independent inductions. *$p<0.05$, ** $p<0.01$, **** $p<0.0001$, one way ANOVA and Tukey's multiple comparison test for multi-group comparisons or t- test for 2-group comparisons **B)** or Kruskal-Wallis test followed by Dunn's multiple comparison test **D)**.

### Supplemental Figure VIII

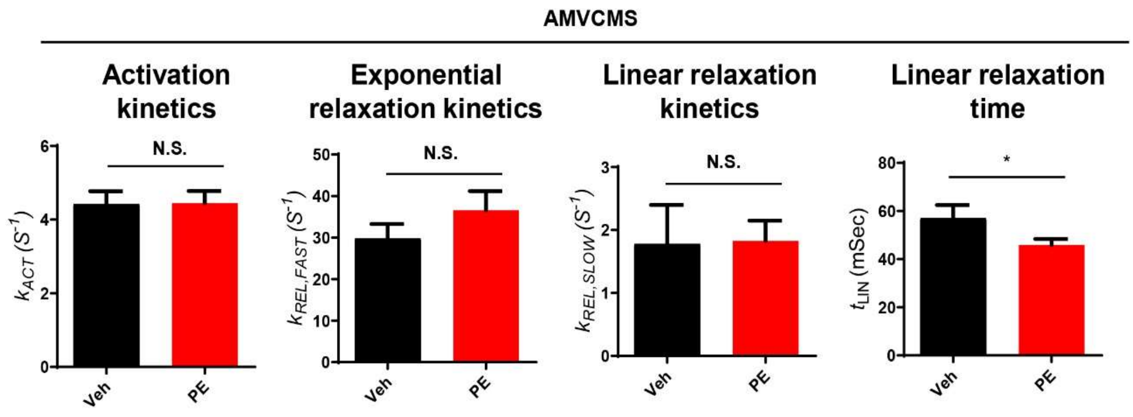


**Figure S8. Myofibril mechanics data on AMVCMs.** Myofibril mechanics data from AMVCMs isolated from 8-12 week old adult mice and treated with vehicle or 10 µmol/L PE for 72 hours before myofibril isolation. Note that rodent myofibrils display profoundly faster kinetics of both activation and relaxation at baseline compared to myofibrils from human hiPSC-CMs and cardiac tissue. *, $p<0.05$, Student *t* test on log2 transformed data. 21-26 myofibrils were isolated from three separate cardiomyocyte isolations. A complete table of myofibril mechanics data on AMVCMs can be seen in **Table S6**.

**Table S1: Myofibril mechanics data.** *, p<0.05 vs GLUC‡, p<0.05 vs donor tissue: one way ANOVA followed by Tukey's multiple comparison test on log2 transformed data.

| | GLUC | MM | MPAT | DONOR HRT |
|---|---|---|---|---|
| **$k_{REL,SLOW}$ ($s^{-1}$)** | 0.20±0.05 | 0.50±0.14 | 0.44±0.10 | 0.48±0.13 |
| **$t_{LIN}$ (ms)** | 144.7±11.2 | 139.6±16.6 | 145.1±13.7 | 155.4±14.0 |
| **$k_{REL,FAST}$ ($s^{-1}$)** | 8.88±0.83 | 9.03±1.21 | 9.51±1.30 | 8.49±1.51 |
| **$k_{ACT}$ ($s^{-1}$)** | 1.04±0.08 | 0.95±0.10 | 1.07±0.13 | 0.84+0.14 |
| **$k_{TR}$ ($s^{-1}$)** | 0.87±0.06 | 0.98±0.07 | 0.76±0.05 | 0.61+0.15 |
| **Max. tens. ($mN/mm^2$)** | 21.40±2.29‡ | 33.74±5.49‡ | 52.10±7.39* | 55.56±5.95 |
| **Rest. tens. ($mN/mm^2$)** | 3.53±0.4‡ | 6.49±1.3 | 6.97±1.13 | 12.22+2.83 |
| **Diameter (µm)** | 4.56±0.22 | 4.00±0.33 | 4.15±0.36 | 4.66±0.50 |
| **N (# Myofibrils)** | 30 | 24 | 31 | 6 |
| **Age** | 32 (age of hiPSC donor) | | | 35 |

**Table S2: Gene ontology on differentially expressed genes.** RNA-seq data was filtered to include only top 12,500 expressed genes for all groups, then GO analysis was performed on genes displaying >1.5 fold regulation between designated groups.

**↑ MPAT vs MM**

| GO Term | Fold Enrichment | P value | FDR |
|---|---|---|---|
| regulation of multicellular organismal process | 1.94 | 5.88E-08 | 9.34E-04 |

**↓ MPAT vs MM**

| GO Term | Fold Enrichment | P value | FDR |
|---|---|---|---|
| cell cycle | 4.08 | 5.00E-32 | 7.95E-28 |
| cell division | 6.67 | 3.26E-29 | 2.59E-25 |
| mitotic cell cycle | 5.3 | 2.43E-27 | 9.64E-24 |
| nuclear division | 7.81 | 1.09E-21 | 2.88E-18 |
| chromosome segregation | 7.77 | 4.23E-21 | 9.61E-18 |
| mitotic nuclear division | 11.4 | 2.40E-20 | 4.24E-17 |
| organelle fission | 7.07 | 2.56E-20 | 4.07E-17 |
| sister chromatid segregation | 10.93 | 1.32E-18 | 1.90E-15 |
| M phase | 9.20 | 1.61E-17 | 2.14E-14 |
| regulation of cell cycle | 3.19 | 4.21E-17 | 4.78E-14 |
| mitotic sister chromatid segregation | 12.23 | 5.76E-17 | 6.11E-14 |

| | | | |
|---|---|---|---|
| microtubule cytoskeleton organization involved in mitosis | 12.12 | 6.90E-17 | 6.86E-14 |
| spindle organization | 9.24 | 1.20E-14 | 1.12E-11 |
| biological phase | 6.27 | 1.30E-14 | 1.15E-11 |
| mitotic cell cycle phase | 6.27 | 1.30E-14 | 1.09E-11 |
| anaphase | 8.59 | 1.33E-14 | 1.00E-11 |
| chromosome organization | 3.12 | 2.92E-14 | 2.02E-11 |
| mitotic spindle organization | 13.4 | 3.81E-14 | 2.52E-11 |
| regulation of cell cycle process | 3.59 | 4.72E-14 | 3.00E-11 |
| mitotic prometaphase | 7.55 | 5.39E-13 | 3.17E-10 |
| regulation of mitotic nuclear division | 7.51 | 6.02E-13 | 3.42E-10 |
| microtubule-based process | 3.67 | 9.27E-13 | 5.08E-10 |
| regulation of chromosome segregation | 9.82 | 9.66E-13 | 5.12E-10 |
| regulation of mitotic sister chromatid separation | 13.51 | 4.75E-12 | 2.44E-09 |
| regulation of mitotic cell cycle | 3.59 | 6.45E-12 | 3.21E-09 |
| cell cycle | 4.08 | 5.00E-32 | 7.95E-28 |
| cell division | 6.67 | 3.26E-29 | 2.59E-25 |
| mitotic cell cycle | 5.3 | 2.43E-27 | 9.64E-24 |

**↑ MPAT vs GLUC**

| **GO Term** | **Fold Enrichment** | **P value** | **FDR** |
|---|---|---|---|

| | | | |
|---|---|---|---|
| nervous system development | 1.72 | 1.97E-14 | 3.13E-10 |
| system development | 1.44 | 1.20E-12 | 4.76E-09 |
| anatomical structure development | 1.38 | 1.82E-12 | 5.78E-09 |
| developmental process | 1.35 | 7.84E-12 | 2.08E-08 |
| cell differentiation | 1.45 | 7.48E-11 | 1.19E-07 |
| regulation of multicellular organismal development | 1.63 | 4.12E-10 | 5.04E-07 |
| regulation of multicellular organismal process | 1.48 | 4.77E-10 | 5.05E-07 |
| anatomical structure morphogenesis | 1.6 | 7.18E-10 | 7.14E-07 |
| cell adhesion | 1.99 | 1.24E-09 | 1.16E-06 |
| axon development | 2.52 | 3.63E-09 | 2.75E-06 |
| cell morphogenesis involved in differentiation | 2.16 | 5.18E-08 | 2.35E-05 |
| cell projection morphogenesis | 2.19 | 1.42E-07 | 5.25E-05 |
| neuron projection morphogenesis | 2.19 | 1.94E-07 | 7.01E-05 |
| plasma membrane bounded cell projection morphogenesis | 2.17 | 2.24E-07 | 7.93E-05 |
| circulatory system development | 1.8 | 1.15E-06 | 3.15E-04 |
| regulation of localization | 1.4 | 1.29E-06 | 3.41E-04 |
| cardiovascular system development | 2.02 | 2.20E-06 | 5.38E-04 |
| plasma membrane bounded cell projection organization | 1.66 | 3.02E-06 | 7.28E-04 |

| | | | |
|---|---|---|---|
| mesenchyme morphogenesis | 4.85 | 1.38E-05 | 2.37E-03 |
| heart valve development | 4.28 | 2.18E-05 | 3.28E-03 |
| endocardial cushion development | 4.98 | 2.38E-05 | 3.54E-03 |
| heart valve morphogenesis | 4.31 | 7.97E-05 | 9.83E-03 |
| cardiac chamber morphogenesis | 2.79 | 1.57E-04 | 1.67E-02 |
| regulation of heart morphogenesis | 4.67 | 1.78E-04 | 1.79E-02 |
| mesenchymal cell differentiation | 2.6 | 1.81E-04 | 1.81E-02 |
| regulation of cholesterol biosynthetic process | 4.34 | 2.94E-04 | 2.55E-02 |
| regulation of lipid metabolic process | 1.87 | 3.13E-04 | 2.60E-02 |
| cardiac chamber development | 2.42 | 3.63E-04 | 2.93E-02 |

**↓ MPAT vs GLUC**

| **GO Term** | **Fold Enrichment** | **P value** | **FDR** |
|---|---|---|---|
| cell cycle | 2.95 | 2.09E-41 | 3.33E-37 |
| mitotic cell cycle | 3.69 | 1.01E-35 | 5.34E-32 |
| cell division | 3.92 | 8.51E-29 | 2.71E-25 |
| nuclear division | 4.54 | 4.18E-22 | 1.11E-18 |
| organelle fission | 4.17 | 1.12E-20 | 2.55E-17 |
| chromosome segregation | 4.23 | 5.98E-19 | 1.19E-15 |
| DNA replication | 4.81 | 9.35E-19 | 1.49E-15 |
| chromosome organization | 2.38 | 9.44E-19 | 1.37E-15 |

| | | | |
|---|---|---|---|
| mitotic nuclear division | 5.9 | 1.16E-18 | 1.54E-15 |
| regulation of cell cycle | 2.26 | 2.36E-18 | 2.89E-15 |
| sister chromatid segregation | 5.8 | 1.15E-17 | 1.30E-14 |
| cytoskeleton organization | 2.26 | 8.07E-17 | 8.55E-14 |
| mitotic prometaphase | 5.04 | 8.12E-17 | 8.07E-14 |
| microtubule cytoskeleton organization | 3.07 | 3.35E-16 | 2.96E-13 |
| biological phase | 3.91 | 6.32E-16 | 5.29E-13 |
| mitotic cell cycle phase | 3.91 | 6.32E-16 | 5.02E-13 |
| organelle organization | 1.6 | 9.17E-16 | 6.34E-13 |
| cellular component organization | 1.42 | 1.35E-15 | 8.93E-13 |
| cell cycle phase transition | 3.73 | 2.12E-15 | 1.35E-12 |
| mitotic cell cycle phase transition | 3.75 | 2.86E-15 | 1.75E-12 |
| DNA conformation change | 3.65 | 4.65E-15 | 2.74E-12 |
| regulation of mitotic cell cycle | 2.57 | 4.62E-14 | 2.53E-11 |
| cellular component organization or biogenesis | 1.38 | 6.39E-14 | 3.39E-11 |
| microtubule-based process | 2.51 | 1.18E-13 | 6.05E-11 |
| DNA-dependent DNA replication | 5.17 | 1.06E-12 | 5.12E-10 |
| M phase (GO:0000279) | 4.27 | 1.57E-12 | 7.36E-10 |
| cell cycle | 2.95 | 2.09E-41 | 3.33E-37 |
| mitotic cell cycle | 3.69 | 1.01E-35 | 5.34E-32 |

**↑ MM vs GLUC**

| GO Term | Fold Enrichment | P value | FDR |
|---|---|---|---|
| anatomical structure development | 1.49 | 4.32E-17 | 6.87E-13 |
| system development | 1.55 | 2.28E-16 | 1.81E-12 |
| cell differentiation | 1.55 | 1.37E-13 | 3.11E-10 |
| regulation of multicellular organismal development | 1.81 | 1.67E-13 | 3.32E-10 |
| cellular developmental process | 1.54 | 1.70E-13 | 3.00E-10 |
| cell adhesion | 2.29 | 3.47E-13 | 5.01E-10 |
| biological adhesion | 2.27 | 6.46E-13 | 8.56E-10 |
| anatomical structure morphogenesis | 1.74 | 2.69E-12 | 3.28E-09 |
| cell projection morphogenesis | 2.41 | 9.34E-09 | 5.30E-06 |
| circulatory system development | 1.97 | 3.56E-08 | 1.38E-05 |
| positive regulation of cell differentiation | 1.89 | 7.77E-08 | 2.69E-05 |
| cell projection organization | 1.78 | 1.63E-07 | 4.55E-05 |
| cardiovascular system development | 2.18 | 4.20E-07 | 8.79E-05 |
| regulation of cell projection organization | 2.01 | 5.68E-07 | 1.13E-04 |
| muscle structure development | 2.17 | 1.89E-06 | 3.22E-04 |
| regulation of lipid metabolic process | 2.18 | 9.06E-06 | 1.29E-03 |
| regulation of cholesterol metabolic process | 4.93 | 1.05E-05 | 1.43E-03 |
| muscle organ development | 2.38 | 1.84E-05 | 2.31E-03 |
| lipid metabolic process | 1.61 | 2.11E-05 | 2.58E-03 |

| | | | |
|---|---|---|---|
| response to glucocorticoid | 3.02 | 2.43E-05 | 2.84E-03 |
| response to growth factor | 1.96 | 2.50E-05 | 2.90E-03 |
| muscle tissue development | 2.28 | 4.34E-05 | 4.72E-03 |
| regulation of cholesterol biosynthetic process | 4.94 | 1.08E-04 | 9.78E-03 |
| regulation of sterol biosynthetic process | 4.94 | 1.08E-04 | 9.72E-03 |
| regulation of steroid metabolic process | 3.09 | 1.09E-04 | 9.75E-03 |
| striated muscle tissue development | 2.24 | 1.54E-04 | 1.27E-02 |
| cardiocyte differentiation | 2.93 | 2.91E-04 | 2.11E-02 |
| regulation of heart morphogenesis | 4.78 | 2.93E-04 | 2.12E-02 |

**↓ MM vs GLUC**

| GO Term | P value | FDR |
|---|---|---|
| Cell cycle | 0.000033242 | 0.0052688 |
| Oocyte meiosis | 0.000033242 | 0.0052688 |
| Progesterone-mediated oocyte maturation | 0.00043059 | 0.045499 |
| ECM-receptor interaction | 0.00098342 | 0.077936 |
| Glycerolipid metabolism | 0.0014225 | 0.090184 |
| Glycine, serine and threonine metabolism | 0.0017477 | 0.092338 |
| cGMP-PKG signaling pathway | 0.0024853 | 0.11255 |
| Calcium signaling pathway | 0.0033015 | 0.13082 |

| | | |
|---|---|---|
| Renin secretion | 0.0080179 | 0.28054 |
| Homologous recombination | 0.0091446 | 0.28054 |

**Table S3: KEGG pathway analysis on differentially expressed genes.** RNA-seq data was filtered to include only top 12,500 expressed genes for all groups, then KEGG Pathway analysis was performed on genes displaying >1.5 fold regulation between designated groups.

**↑ MPAT vs MM**

| KEGG Term | P value | FDR |
|---|---|---|
| Glycine, serine and threonine metabolism | 0.001581 | 0.50119 |
| ErbB signaling pathway | 0.0044073 | 0.53317 |
| Salivary secretion | 0.0056192 | 0.53317 |
| Axon guidance | 0.0067277 | 0.53317 |
| Adrenergic signaling in cardiomyocytes | 0.009867 | 0.62557 |
| PPAR signaling pathway | 0.014347 | 0.75802 |
| Phenylalanine metabolism | 0.01923 | 0.76198 |
| Selenocompound metabolism | 0.01923 | 0.76198 |
| Mineral absorption | 0.026744 | 0.94197 |
| Phosphatidylinositol signaling system | 0.036996 | 1 |

**↓ MPAT vs MM**

| KEGG Term | P value | FDR |
|---|---|---|
| Cell cycle | 1.53E-11 | 4.86E-09 |

| | | |
|---|---|---|
| Human T-cell leukemia virus 1 infection | 2.78E-08 | 4.4127E-06 |
| Cellular senescence | 9.2739E-06 | 0.00097994 |
| p53 signaling pathway | 0.00092778 | 0.067806 |
| Hepatitis B | 0.0011578 | 0.067806 |
| Breast cancer | 0.0013546 | 0.067806 |
| Oocyte meiosis | 0.0014973 | 0.067806 |
| MAPK signaling pathway | 0.0019064 | 0.075543 |
| Colorectal cancer | 0.0026241 | 0.092425 |
| Small cell lung cancer | 0.0040755 | 0.12394 |

↑ **MPAT vs GLUC**

| KEGG Term | P value | FDR |
|---|---|---|
| Fatty acid metabolism | 0.000010053 | 0.003187 |
| Fatty acid degradation | 0.00013291 | 0.021066 |
| Axon guidance | 0.00033531 | 0.035431 |
| Metabolic pathways | 0.00055519 | 0.043999 |
| AGE-RAGE signaling pathway in diabetic complications | 0.0012805 | 0.081183 |
| Protein processing in endoplasmic reticulum | 0.0023937 | 0.1258 |
| PPAR signaling pathway | 0.0027779 | 0.1258 |
| Glycosaminoglycan biosynthesis | 0.0043429 | 0.17209 |
| Fluid shear stress and atherosclerosis | 0.005151 | 0.18138 |

| | | |
|---|---|---|
| Fatty acid elongation | 0.0059162 | 0.181 |

**↓ MPAT vs GLUC**

| KEGG Term | P value | FDR |
|---|---|---|
| Cell cycle | 7.05E-13 | 2.24E-10 |
| Oocyte meiosis | 1.1788E-06 | 0.00018683 |
| DNA replication | 2.5478E-06 | 0.00026921 |
| Fanconi anemia pathway | 0.00003304 | 0.0023315 |
| p53 signaling pathway | 0.000036774 | 0.0023315 |
| Homologous recombination | 0.000064576 | 0.0034118 |
| Progesterone-mediated oocyte maturation | 0.00011021 | 0.0049911 |
| Cellular senescence | 0.0014981 | 0.059361 |
| Small cell lung cancer | 0.0019308 | 0.063087 |
| Base excision repair | 0.0019901 | 0.063087 |

**↑ MM vs GLUC**

| KEGG Term | P value | FDR |
|---|---|---|
| Fatty acid metabolism | 0.00017398 | 0.044812 |
| Carbon metabolism | 0.00040047 | 0.044812 |
| Fatty acid degradation | 0.00042409 | 0.044812 |
| Metabolic pathways | 0.00083683 | 0.066319 |
| Fluid shear stress and atherosclerosis | 0.0010884 | 0.069008 |
| Cell adhesion molecules (CAMs) | 0.001613 | 0.074791 |

| | | |
|---|---|---|
| Biosynthesis of amino acids | 0.0019203 | 0.074791 |
| MAPK signaling pathway | 0.00201 | 0.074791 |
| Alanine, aspartate and glutamate metabolism | 0.0021234 | 0.074791 |
| Steroid biosynthesis | 0.0026163 | 0.082938 |

**↓ MM vs GLUC**

| KEGG Term | P value | FDR |
|---|---|---|
| Cell cycle | 0.000033242 | 0.0052688 |
| Oocyte meiosis | 0.000033242 | 0.0052688 |
| Progesterone-mediated oocyte maturation | 0.00043059 | 0.045499 |
| ECM-receptor interaction | 0.00098342 | 0.077936 |
| Glycerolipid metabolism | 0.0014225 | 0.090184 |
| Glycine, serine and threonine metabolism | 0.0017477 | 0.092338 |
| cGMP-PKG signaling pathway | 0.0024853 | 0.11255 |
| Calcium signaling pathway | 0.0033015 | 0.13082 |
| Renin secretion | 0.0080179 | 0.28054 |
| Homologous recombination | 0.0091446 | 0.28054 |

**Tables S4-6: Myofibril mechanics data.** *, **, **** $p<0.05$, 0.01, 0.0001 vs PBS-treated (Table 2 and 4) or vs Donor (Table 3), t-test on log2 transformed data.

**Table S4:** PE-treated hiPSC-CM Myofibril Characteristics

| | **hiPSC-CM+Veh** | **hiPSC-CM+PE** |
|---|---|---|
| **$k_{REL,SLOW}$ ($s^{-1}$)** | 0.38±0.07 | 0.58±0.12 |
| **$t_{LIN}$ (ms)** | 196.0±17.4 | 113.6±6.2**** |
| **$k_{REL,FAST}$ ($s^{-1}$)** | 6.23±0.63 | 8.64±0.77* |
| **$k_{ACT}$ ($s^{-1}$)** | 0.88±0.09 | 1.03±0.08 |
| **$k_{TR}$ ($s^{-1}$)** | 0.76±0.04 | 0.80±0.06 |
| **Max. tens. ($mN/mm^2$)** | 47.96±5.91 | 44.04±5.05 |
| **Rest. tens. ($mN/mm^2$)** | 7.06±1.16 | 6.15±0.90 |
| **Diameter (μm)** | 4.49±0.26 | 4.48±0.24 |
| **N (# myofibrils)** | 45 | 44 |

**Table S5:** Human Donor and HCM Myofibril Characteristics

| | Donor HRT | HCM HRT |
|---|---|---|
| $k_{REL,SLOW}$ ($s^{-1}$) | 0.45±0.09 | 0.83±0.25 |
| $t_{LIN}$ (ms) | 157.1±3.2 | 114.6±15.3* |
| $k_{REL,FAST}$ ($s^{-1}$) | 8.53±0.50 | 10.82±3.19 |
| $k_{ACT}$ ($s^{-1}$) | 0.78±0.16 | 1.38±0.17* |
| $k_{TR}$ ($s^{-1}$) | 0.53±0.04 | 0.74±0.05** |
| Max. tens. ($mN/mm^2$) | 53.55±12.61 | 47.81±5.03 |
| Rest. tens. ($mN/mm^2$) | 11.50±2.83 | 8.03±2.33 |
| Diameter (µm) | 5.62±0.37 | 4.43±0.15* |
| N (# hrts) | 4 | 4 |
| Age | 31.25±0.48 | 47.5±3.92* |
| Sex (#male, # female) | 4:0 | 3:1 |

**Table S6:** PE-treated AMVCM Myofibril Characteristics

| | AMVCM+PBS | AMVCM+PE |
|---|---|---|
| $k_{REL,SLOW}$ ($s^{-1}$) | 1.78±0.62 | 1.83±0.32 |
| $t_{LIN}$ (ms) | 57.15±5.36 | 45.83±2.58* |
| $k_{REL,FAST}$ ($s^{-1}$) | 29.79±3.46 | 36.61±4.59 |
| $k_{ACT}$ ($s^{-1}$) | 4.42±0.35 | 4.45±0.33 |
| $k_{TR}$ ($s^{-1}$) | 3.57±0.30 | 3.12±0.30 |
| Max. tens. (mN/mm$^2$) | 57.52±5.04 | 68.55±6.39 |
| Rest. tens. (mN/mm$^2$) | 4.72±0.71 | 6.94±1.07 |
| Diameter (μm) | 4.66±0.29 | 4.12±0.22 |
| N (# myofibrils) | 21 | 26 |

Supplementary Video 1. Spontaneous contraction of hiPSC-CMs spread sheet under the GLUC condition.

Supplementary Video 2. Spontaneous contraction of hiPSC-CMs spread sheet under the MM condition.

Supplementary Video 3. Spontaneous contraction of hiPSC-CMs spread sheet under the MPAT condition.

### References


5. Lian X, Hsiao C, Wilson G, Zhu K, Hazeltine LB, Azarin SM, Raval KK, Zhang J, Kamp TJ and Palecek SP. Robust cardiomyocyte differentiation from human pluripotent stem cells via temporal modulation of canonical Wnt signaling. *Proc Natl Acad Sci U S A*. 2012;109:E1848-57.

6. Tohyama S, Hattori F, Sano M, Hishiki T, Nagahata Y, Matsuura T, Hashimoto H, Suzuki T, Yamashita H, Satoh Y, Egashira T, Seki T, Muraoka N, Yamakawa H, Ohgino Y, Tanaka T, Yoichi M, Yuasa S, Murata M, Suematsu M and Fukuda K. Distinct metabolic flow enables large-scale purification of mouse and human pluripotent stem cell-derived cardiomyocytes. *Cell Stem Cell*. 2013;12:127-37.

40. Nemkov T, Reisz JA, Gehrke S, Hansen KC and D'Alessandro A. High-Throughput Metabolomics: Isocratic and Gradient Mass Spectrometry-Based Methods. *Methods Mol Biol*. 2019;1978:13-26.

41. Nemkov T, Hansen KC and D'Alessandro A. A three-minute method for high-throughput quantitative metabolomics and quantitative tracing experiments of central carbon and nitrogen pathways. *Rapid Commun Mass Spectrom*. 2017;31:663-673.

42. Sparagna GC, Johnson CA, McCune SA, Moore RL and Murphy RC. Quantitation of cardiolipin molecular species in spontaneously hypertensive heart failure rats using electrospray ionization mass spectrometry. *J Lipid Res*. 2005;46:1196-204.

43. Bligh EG and Dyer WJ. A rapid method of total lipid extraction and purification. *Can J Biochem Physiol*. 1959;37:911-917.

44. Yoon HR, Hong YM, Boriack RL and Bennett MJ. Effect of L-carnitine supplementation on cardiac carnitine palmitoyltransferase activities and plasma carnitine concentrations in adriamycin-treated rats. *Pediatr Res*. 2003;53:788-92.

45. FastQC [Internet]. 2019.

46. Dobin A, Davis CA, Schlesinger F, Drenkow J, Zaleski C, Jha S, Batut P, Chaisson M and Gingeras TR. STAR: ultrafast universal RNA-seq aligner. *Bioinformatics*. 2013;29:15-21.

47. Liao Y, Smyth GK and Shi W. featureCounts: an efficient general purpose program for assigning sequence reads to genomic features. *Bioinformatics*. 2014;30:923-30.

48. Liao Y, Smyth GK and Shi W. The R package Rsubread is easier, faster, cheaper and better for alignment and quantification of RNA sequencing reads. *Nucleic Acids Res*. 2019;47:e47.

49. Love MI, Huber W and Anders S. Moderated estimation of fold change and dispersion for RNA-seq data with DESeq2. *Genome Biol*. 2014;15:550.

50. Mi H, Muruganujan A, Ebert D, Huang X and Thomas PD. PANTHER version 14: more genomes, a new PANTHER GO-slim and improvements in enrichment analysis tools. *Nucleic Acids Res*. 2019;47:D419-D426.
